# Insulin-like growth factor-1 (IGF-1), insulin-like growth factor-binding protein-3 (IGFBP-3) and breast cancer risk: observational and Mendelian randomization analyses

**DOI:** 10.1101/809046

**Authors:** Neil Murphy, Anika Knuppel, Nikos Papadimitriou, Richard M Martin, Konstantinos K Tsilidis, Karl Smith-Byrne, Georgina Fensom, Aurora Perez-Cornago, Ruth C Travis, Timothy J Key, Marc J Gunter

## Abstract

**Background:** Epidemiological evidence supports a positive association between circulating insulin-like growth factor-1 (IGF-1) concentrations and breast cancer risk, but both the magnitude and causality of this relationship are uncertain. We conducted observational analyses with adjustment for regression dilution bias, and Mendelian randomization (MR) analyses to allow causal inference.

**Patients and methods:** We investigated the associations between circulating IGF-1 concentrations and incident breast cancer risk in 206,263 women in the UK Biobank. Multivariable hazard ratios (HRs) and 95% confidence intervals (CI) were estimated using Cox proportional hazards models. HRs were corrected for regression dilution using repeat IGF-1 measures available in a subsample of 6,711 women. For the MR analyses, genetic variants associated with circulating IGF-1 and IGFBP-3 levels were identified and their association with breast cancer was examined with two-sample MR methods using genome-wide data from 122,977 cases and 105,974 controls.

**Results:** In the UK Biobank, after a median follow-up of 7.1 years, 4,360 incident breast cancer cases occurred. In the multivariable-adjusted models corrected for regression dilution, higher IGF-1 concentrations were associated with a greater risk of breast cancer (HR per 5 nmol/L increment of IGF-1=1.11, 95%CI=1.07-1.16). Similar positive associations were found by follow-up time, menopausal status, body mass index, and other risk factors. In the MR analyses, a 5 nmol/L increment in genetically-predicted IGF-1 concentration was associated with greater breast cancer risk (odds ratio [OR]=1.05, 95%CI=1.01-1.10; Pvalue=0.02), with a similar effect estimate for estrogen positive (ER+) tumors, but no effect found for estrogen negative (ER^-^) tumors. Genetically-predicted IGFBP-3 concentrations were not associated with breast cancer risk (OR per 1-SD increment=1.00, 95%CI=0.97-1.04; Pvalue=0.98).

**Conclusion:** Our results support a probable causal relationship between circulating IGF-1 concentrations and breast cancer, suggesting that interventions targeting the IGF pathway may be beneficial in preventing breast tumorigenesis.

**Disclaimer:** Where authors are identified as personnel of the International Agency for Research on Cancer / World Health Organization, the authors alone are responsible for the views expressed in this article and they do not necessarily represent the decisions, policy or views of the International Agency for Research on Cancer / World Health Organization.

## Introduction

Insulin-like growth factor-1 (IGF-1) is a polypeptide that has mitogenic and anti-apoptotic effects^1,2^. Approximately 99% of IGF-1 is bound to IGF binding proteins, with most bound to insulin-like growth factor-binding protein-3 (IGFBP-3)^3^. In experimental studies, IGFBP-3 has also been shown to not only regulate IGF-1 bioavailability, but also to have direct inhibitory effects on cell growth^4^.

Interest in the possible role of IGF-1 in the development of breast cancer began in the 1980s^5^. An early case-control study reported higher plasma concentrations of IGF-1 in women with breast cancer than in controls^6^, and in the first prospective study circulating concentrations of IGF-1 were positively associated with breast cancer risk for premenopausal women, but not postmenopausal women^7^. Most subsequent prospective studies have supported a positive association between IGF-1 and breast cancer risk, and a pooled individual-participant data analysis of 4,790 cases from 17 prospective studies showed that women with relatively high circulating IGF-1 had a ∼30% higher risk of breast cancer than women with relatively low circulating IGF-1; there was no evidence that the association was due to reverse-causation, varied by menopausal status, or was attenuated by adjustment for other risk factors including IGFBP-3, reproductive factors, and body mass index (BMI)^8^. However, heterogeneity by estrogen receptor (ER) subtype was found, with the positive association present for ER^+^, but not ER^-^ tumours. In addition, this pooled analysis was based on a single IGF-1 measure for each woman so risk estimates would have been influenced by the combined effects of measurement error and within-person variability, leading to a likely underestimation of the true association (regression dilution)^8,9^.

To further examine the possible causal role of IGF-1 in breast cancer risk, we conducted complementary observational and Mendelian randomization (MR) analyses. Firstly, we investigated how prediagnostic circulating concentrations of IGF-1 were related to breast cancer risk in the UK Biobank study, a large prospective cohort in which a subsample of participants have repeat IGF-1 measures enabling correction for regression dilution bias. Next, we used a two-sample MR approach to examine potential causal associations by combining genetic variants associated with circulating IGF-1 and IGFBP-3 concentrations in genome-wide association studies (GWAS), and then assessing the association of these variants with breast cancer (overall, ER^+^, and ER^-^) risk in a large consortium of 122,977 breast cancer cases and 105,974 controls^10^.

## Methods

### UK Biobank – observational analysis

#### Study participants

The UK Biobank is a prospective cohort of 502,536 adults aged between 40 and 69 years (229,182 men and 273,474 women) who were recruited between 2006 and 2010^11^. The UK Biobank invited ∼9.2 million people to participate through postal invitation with a telephone follow-up, with a response rate of 5.7%. All participants were registered with the UK National Health Service (NHS) and lived within ∼25 miles (40 km) of one of the 22 study assessment centres. The UK Biobank has approval from the North West Multi-centre Research Ethics Committee, the National Information Governance Board for Health and Social Care in England and Wales, and the Community Health Index Advisory Group in Scotland. In addition, an independent Ethics and Governance Council was formed in 2004 to oversee UK Biobank’s continuous adherence to the Ethics and Governance Framework which was developed for the study (http://www.ukbiobank.ac.uk/ethics/). All participants provided written informed consent at recruitment and to be followed up using data-linkage. This research has been conducted using the UK Biobank Resource under application numbers 3248 and 24494.

During the baseline recruitment visit, participants were asked to complete a self-administered touchscreen questionnaire, which included questions on socio-demographics (including age, sex, education and postcode, used to assign Townsend deprivation score), health/medical history, and lifestyle exposures (including smoking habits, dietary intakes, and alcohol consumption). At the baseline visit, participants also underwent physical measurements, including body weight, height, and waist circumference. Blood samples were collected from all participants at recruitment, and repeat blood samples were collected from a subset of ∼20,000 participants who re-attended the assessment center between 2012 and 2013. Blood samples were centrifuged, and serum stored at −80°C.

Exclusions prior to the onset of analyses were men (N=229,134); women with prevalent cancer (including *in situ* breast cancer, but excluding non-malignant skin cancer) at recruitment (N=18,654); participants in whom genetic sex differed from reported gender (N=121), missing data on body size measurements (N=1,350); prevalent type-2 diabetes or unknown diabetes status at recruitment based on hospital records and self-report (because diabetes medications can affect circulating concentrations of IGF-1^12^; N=10,705); women who reported oral contraceptive and menopausal hormone use at recruitment (because oral oestrogens alter hepatic protein production and change circulating concentrations of IGF-1^12^; N=20,988); and participants without an IGF-1 measurement (N=15,415). Our analysis therefore included 206,263 women.

#### Blood collection and laboratory methods

As part of the UK Biobank Biomarker Project^13^, serum concentrations of IGF-1 (DiaSorin Liaison XL), estradiol, testosterone, and sex hormone binding globulin (SHBG) were determined by a chemiluminescent immunoassay (Beckman Coulter DXI 800). The Immuno-turbidimetric method (Beckman Coulter DXI 800) was used to assay serum high sensitivity C-reactive protein (CRP) concentrations. Glycated haemoglobin (HbA1c) concentrations were determined using the HPLC Variant II Turbo 2.0 system (Bio–Rad). Full details on assay performance have been published^13^. In summary, average within-laboratory (total) coefficient of variation (CV) for low, medium, and high internal quality control level samples for each biomarker ranged from 1.7-15.3% (for IGF-1, the CVs ranged from 5.3-6.2%)^13^. A total of 6,711 women had IGF-1 concentrations measured in blood samples collected at both the recruitment and repeat assessment visit (median of 4-years apart).

#### Assessment of outcome

Incident cancer cases and cancer cases first recorded in death certificates within the UK Biobank cohort were identified through linkage to national cancer registries and death records. Complete follow-up was available until 31st March 2016 for England and Wales and 31st October 2015 for Scotland. Cancer incidence data were coded using the 10th Revision of the International Classification of Diseases (ICD-10). Breast cancer was defined as registration ICD-10: C50.

#### Statistical analysis

Hazard ratios (HRs) and 95% confidence intervals (CIs) were estimated using Cox proportional hazards models. Age was the primary time variable in all models. Time at entry was age at recruitment. Exit time was age at whichever of the following came first: breast cancer diagnosis, breast cancer at death without prior diagnosis, death, or the last date at which follow-up was considered complete. Models were stratified by age at recruitment in 5-year categories, Townsend deprivation index fifths, and region of the recruitment assessment centre. Deviations from proportionality were assessed using an analysis of Schoenfeld residuals, with no evidence of non-proportionality being detected. IGF-1 was modelled on the continuous scale (per 5 nmol/L) and with participants grouped into sex-specific fifths of circulating concentrations.

HRs were additionally corrected for regression dilution using regression dilution ratios obtained from the subsample of 6,711 women with repeated IGF-1 measurement^9,14^; to obtain corrected HRs the log HRs and their standard errors were divided by the regression dilution ratio for IGF-1 (0.74) and then exponentiated^15^. Possible nonlinear effects were modelled using restricted cubic spline models with five knots placed at Harrell’s default percentiles of circulating IGF-1 concentrations^16^.

The multivariable model (model 1) was adjusted for a set of breast cancer risk factors determined *a priori* namely total physical activity, height, alcohol consumption frequency, smoking status and intensity, educational level, ever use of hormone replacement therapy (HRT), parity and age at first birth, and an interaction between menopausal status and BMI. We also additionally adjusted the multivariable models (model 2) for markers of inflammatory, sex hormone, and glycemic pathways that are known to interrelate/crosstalk with the IGF system^12^, namely CRP, testosterone, SHBG, and HbA1c. Statistical tests for trend were calculated using the ordinal fifths of IGF-1 entered into the model as a continuous variable.

The circulating IGF-1 and breast cancer associations were further assessed across subgroups of BMI, height, menopausal status at recruitment, ages at blood collection and diagnosis, follow-up time, smoking status, and circulating concentrations of CRP, HbA1c, testosterone, and SHBG. Interaction terms (multiplicative scale) between these variables and circulating IGF-1 concentrations were included in separate models, and the statistical significance of the cross-product terms were evaluated using the likelihood ratio test, or competing risk for follow-up time and age at diagnosis.

Statistical tests were all two-sided and a Pvalue<0.05 was considered statistically significant. Analyses were conducted using Stata version 14.

### Mendelian randomization

#### Genetic determinants of insulin-like growth factor 1 (IGF-1) and insulin-like growth factor-binding protein-3 (IGFBP-3)

Genetic markers for circulating IGF-1 and IGFBP-3 concentrations comprised SNPs identified (Pvalue<5 × 10^−8^) from the largest GWAS to date^17,18^. For IGF-1, this GWAS was of 194,174 women from the UK Biobank^18^. The GWAS analyses of IGFBP-3 combined data on 18,995 individuals (men and women) from 13 studies^17^. All participants were of European ancestry. From the genome-wide significant variants identified in these GWAS, we excluded correlated SNPs based on a linkage disequilibrium (LD) level of R^2^<0.01. The instruments for IGF-1 (265 SNPs) and IGFBP-3 (4 SNPs) explained 5.2% and 6.1% of variability in circulating concentrations, respectively. Summary information on the genetic instruments, and the effect estimates for each individual SNP with IGF-1 and IGFBP-3 concentrations, are presented in Tables S1 and S2.

#### Data on breast cancer

Summary data for the associations of the IGF-1 and IGFBP-3 related genetic variants with breast cancer were obtained from a GWAS of 228,951 women (122,977 breast cancer [69,501 ER+, 21,468 ER^-^] cases and 105,974 controls) of European ancestry from the Breast Cancer Association Consortium (BCAC)^10^. Genotypes were imputed using the 1000 Genomes Project reference panel and the regression models adjusted for the first ten principal components and country or study. Effect estimates for the association of each individual SNP with breast cancer are presented in Table S2.

#### Statistical analysis

Two-sample MR analyses using summary data and an inverse variance weighted approach were implemented. MR results correspond to an odds ratio (OR) per 5 nmol/L of genetically-predicted IGF-1 concentration and per 1-standard deviation (SD) of IGFBP-3. Heterogeneity of associations across breast cancer subtypes was assessed by calculating χ^2^ statistics. Cochran’s Q statistics quantified heterogeneity across the individual SNPs. Sensitivity analyses were used to check and correct for the presence of pleiotropy in the causal estimates. To evaluate the extent to which directional pleiotropy (non-balanced horizontal pleiotropy in the MR risk estimates) may have affected the causal estimates for the IGF-1 and breast cancer association, we used an MR-Egger regression approach^19^. We also computed OR estimates using the complementary weighted-median method that can give valid MR estimates under the presence of pleiotropy when up to 50% of the included instruments are invalid^20^. The presence of pleiotropy was also assessed using the MR pleiotropy residual sum and outlier test (MR-PRESSO). In this, outlying SNPs are excluded from the instruments and the effect estimates are reassessed^21^. As a visual evaluation of directional pleiotropy (asymmetry), we also examined a funnel plot of the effect estimate and standard error of each SNP within the IGF-1 instrument on breast cancer risk. For the IGFBP-3 instrument, we conducted leave–one–out analyses to assess the influence of individual variants on the observed associations. All statistical analyses were performed using the *MendelianRandomization* R package^22^.

## Results

### UK Biobank – observational analysis

After a median follow-up time of 7.1 years, 4,360 incident breast cancer cases were recorded. Compared to those in the lowest fifth, participants in the highest circulating IGF-1 fifth were younger, taller, had lower BMI and waist circumference, and were more likely to be never smokers, nulliparous, and never HRT users (Table 1). In addition, participants in the highest circulating IGF-1 fifth had lower circulating concentrations of CRP, HbA1c, and SHBG, with higher circulating concentrations of testosterone.

**Table 1.** Characteristics of UK Biobank study participants by fifth of circulating insulin-like growth factor-1 (IGF-1) concentrations (N=206,263 women)

| Characteristic | Insulin-like growth factor-1 (IGF-1) concentrations |  |  |  |  |
| --- | --- | --- | --- | --- | --- |
|  | 1<br>(N=41,264) | 2<br>(N=41,256) | 3<br>(N=41,239) | 4<br>(N=41,264) | 5<br>(N=41,240) |
| **IGF-1 at baseline (nmol/L)** | 13.8 (2.0) | 18.0 (0.9) | 20.9 (0.8) | 23.8 (0.9) | 29.4 (3.9) |
| **IGF-1 at follow-up (nmol/L), (N=6,711)** | 14.9 (3.5) | 18.0 (3.3) | 20.2 (3.4) | 22.4 (3.6) | 26.4 (5.2) |
| **Breast cancer, N** | 836 | 818 | 911 | 895 | 900 |
| **Age at baseline (years)** | 59.4 (6.9) | 57.8 (7.4) | 56.4 (7.8) | 54.9 (8.0) | 52.6 (8.2) |
| **BMI (kg/m<sup>2</sup>)** | 28.3 (6.0) | 27.2 (5.2) | 26.7 (4.8) | 26.4 (4.5) | 26.0 (4.2) |
| **Waist circumference (cm)** | 87.5 (13.8) | 84.9 (12.4) | 83.7 (11.7) | 82.9 (11.2) | 82.0 (10.6) |
| **Standing height (cm)** | 161.5 (6.3) | 162.1 (6.2) | 162.5 (6.3) | 162.9 (6.3) | 163.5 (6.3) |
| **Socio-economic status (Townsend deprivation index), N (%)** |  |  |  |  |  |
| Most deprived fifth | 9330 (22.6) | 8255 (20.0) | 7799 (18.9) | 7921 (19.2) | 7898 (19.2) |
| **Qualification, N (%)** |  |  |  |  |  |
| Professional qualification, college/university degree | 21109 (51.2) | 22767 (55.2) | 23763 (57.6) | 24643 (59.7) | 25824 (62.6) |
| **Smoking, N (%)** |  |  |  |  |  |
| Never smoker | 23657 (57.3) | 24223 (58.7) | 24830 (60.2) | 25200 (61.1) | 25806 (62.6) |
| Current smoker, ≥15 cigs/day | 1519 (3.7) | 1450 (3.5) | 1310 (3.2) | 1367 (3.3) | 1352 (3.3) |
| **Alcohol intake, N (%)** |  |  |  |  |  |
| Never | 4987 (12.1) | 3712 (9.0) | 3354 (8.1) | 3239 (7.8) | 3128 (7.6) |
| Daily/almost daily | 6852 (16.6) | 7313 (17.7) | 7036 (17.1) | 6666 (16.2) | 5586 (13.5) |
| **Physical activity, N (%)** |  |  |  |  |  |
| <10 METhrs/wk | 10481 (25.4) | 9546 (23.1) | 9314 (22.6) | 9216 (22.3) | 8921 (21.6) |
| ≥60 METhrs/wk | 8245 (20.0) | 8544 (20.7) | 8379 (20.3) | 8383 (20.3) | 8142 (19.7) |
| **HRT use, N (%)** |  |  |  |  |  |
| Never | 23135 (56.1) | 25505 (61.8) | 27310 (66.2) | 28846 (69.9) | 30992 (75.2) |
| Past | 18129 (43.9) | 15751 (38.2) | 13929 (33.8) | 12418 (30.1) | 10248 (24.9) |
| **OCP use, N (%)** |  |  |  |  |  |
| Never | 9642 (23.4) | 8376 (20.3) | 7802 (18.9) | 6896 (16.7) | 6485 (15.7) |
| Past | 31622 (76.6) | 32880 (79.7) | 33437 (81.1) | 34368 (83.3) | 34755 (84.3) |
| **Parity, age at 1st birth N (%)** |  |  |  |  |  |
| Nulliparous | 6650 (16.1) | 6854 (16.6) | 7551 (18.3) | 7816 (18.9) | 8709 (21.1) |
| **Menopausal status, N (%)** |  |  |  |  |  |
| Pre- | 4598 (11.1) | 7382 (17.9) | 9642 (23.4) | 12497 (30.3) | 16675 (40.4) |
| Post- | 35099 (85.1) | 32049 (77.7) | 29551 (71.7) | 26537 (64.3) | 22041 (53.4) |
| **C-reactive protein (mg/L)** | 3.90 (5.6) | 2.77 (4.2) | 2.3 (3.6) | 2.0 (3.4) | 1.65 (3.1) |
| **Glycated hemoglobin (HbA1c; mmol/mol)** | 35.9 (4.9) | 35.4 (4.2) | 35.1 (4.1) | 34.9 (4.0) | 34.6 (3.9) |
| **Testosterone (nmol/L)** | 1.1 (0.6) | 1.1 (0.6) | 1.1 (0.6) | 1.2 (0.6) | 1.2 (0.7) |
| **Sex hormone binding globulin (SHBG; nmol/L)** | 64.5 (32.1) | 62.1 (28.7) | 60.9 (27.3) | 59.7 (26.3) | 57.2 (25.1) |

Mean and standard deviation unless specified. Abbreviations: BMI = body mass index; Cig - cigarettes; METhrs = metabolic equivalent hours; HRT = hormone replacement therapy; OCP = oral contraceptive pill.

#### Association between circulating insulin-like growth factor 1 (IGF-1) concentrations and breast cancer risk

Circulating IGF-1 concentrations were positively associated with breast cancer risk in the minimally adjusted model (HR for highest versus lowest fifth=1.23, 95%CI=1.12–1.36; Ptrend<0.0001) (Table 2). Statistical adjustment for other breast cancer risk factors and circulating concentrations of CRP, HbA1c, testosterone, and SHBG did not materially change the association (HR for highest versus lowest fifth=1.24, 95%CI=1.12–1.37; Ptrend<0.0001). In the restricted cubic spline model, no deviation from linearity for the relationship between IGF-1 and breast cancer was observed (Pnon-linear=0.85). In the continuous multivariable model, adjusted for circulating concentrations of CRP, HbA1c, testosterone, and SHBG, a 5 nmol/L increment in IGF-1 was associated with a higher breast cancer risk (HR=1.08, 95%CI=1.05-1.11). Subsequent correction for regression dilution bias resulted in a larger positive relationship (HR=1.11, 95%CI=1.07-1.15). Similar magnitude positive associations were found according to subgroups of follow-up time, ages at blood collection and diagnosis, menopausal status at recruitment, and other breast cancer risk factors (Pheterogeneities>0.08) (Figure 1, Table S3).

**Table 2.** Risk (hazard ratios) of breast cancer associated with circulating insulin-like growth factor-1 (IGF-1) concentrations in the UK Biobank

| IGF-1 at baseline<br>(nmol/L) | N Cases /<br>Participants | HR (95%CI)<br>Model <sup>0</sup> | HR (95%CI)<br>Model <sup>1</sup> | HR (95%CI)<br>Model <sup>2</sup> |
| --- | --- | --- | --- | --- |
| **Fifths** |  |  |  |  |
| 1 (<16.4) | 836 / 40428 | 1 (reference) | 1 (reference) | 1 (reference) |
| 2 (16.4-<19.5) | 818 / 40438 | 1.00 (0.91-1.10) | 1.01 (0.92-1.11) | 1.01 (0.91-1.11) |
| 3 (19.5-<22.3) | 911 / 40328 | 1.15 (1.05-1.27) | 1.16 (1.06-1.28) | 1.15 (1.05-1.27) |
| 4 (22.3-<25.6) | 895 / 40369 | 1.16 (1.06-1.28) | 1.18 (1.07-1.30) | 1.17 (1.06-1.29) |
| 5 (≥25.6) | 900 / 40340 | 1.23 (1.12-1.36) | 1.25 (1.13-1.38) | 1.24 (1.12-1.37) |
| Ptrend |  | <0.0001 | <0.0001 | <0.0001 |
| **Per 5 nmol/L**<br>**increment** | 4360 / 206263 | 1.08 (1.05-1.11) | 1.08 (1.05-1.11) | 1.08 (1.05-1.11) |
| **Per 5 nmol/L**<br>**increment (adjusted) <sup>a</sup>** | 4360 / 206263 | 1.11 (1.07-1.15) | 1.11 (1.07-1.15) | 1.11 (1.07-1.16) |

HR=hazard ratio; CI=confidence interval; SD=standard deviation.
Model <sup>2</sup>: Model <sup>1</sup> plus additional adjustment for circulating concentrations (fifths, missing/unknown) of C-reactive protein (CRP; mg/L), glycated hemoglobin (HbA1c; mmol/mol), testosterone (nmol/L), and sex hormone binding globulin (SHBG; nmol/L).
<sup>a</sup>: HRs per SD increment were additionally corrected for regression dilution using a regression dilution ratio (0.74) obtained from the subsample of women with repeat IGF-1 measurements.

**Figure 1.**
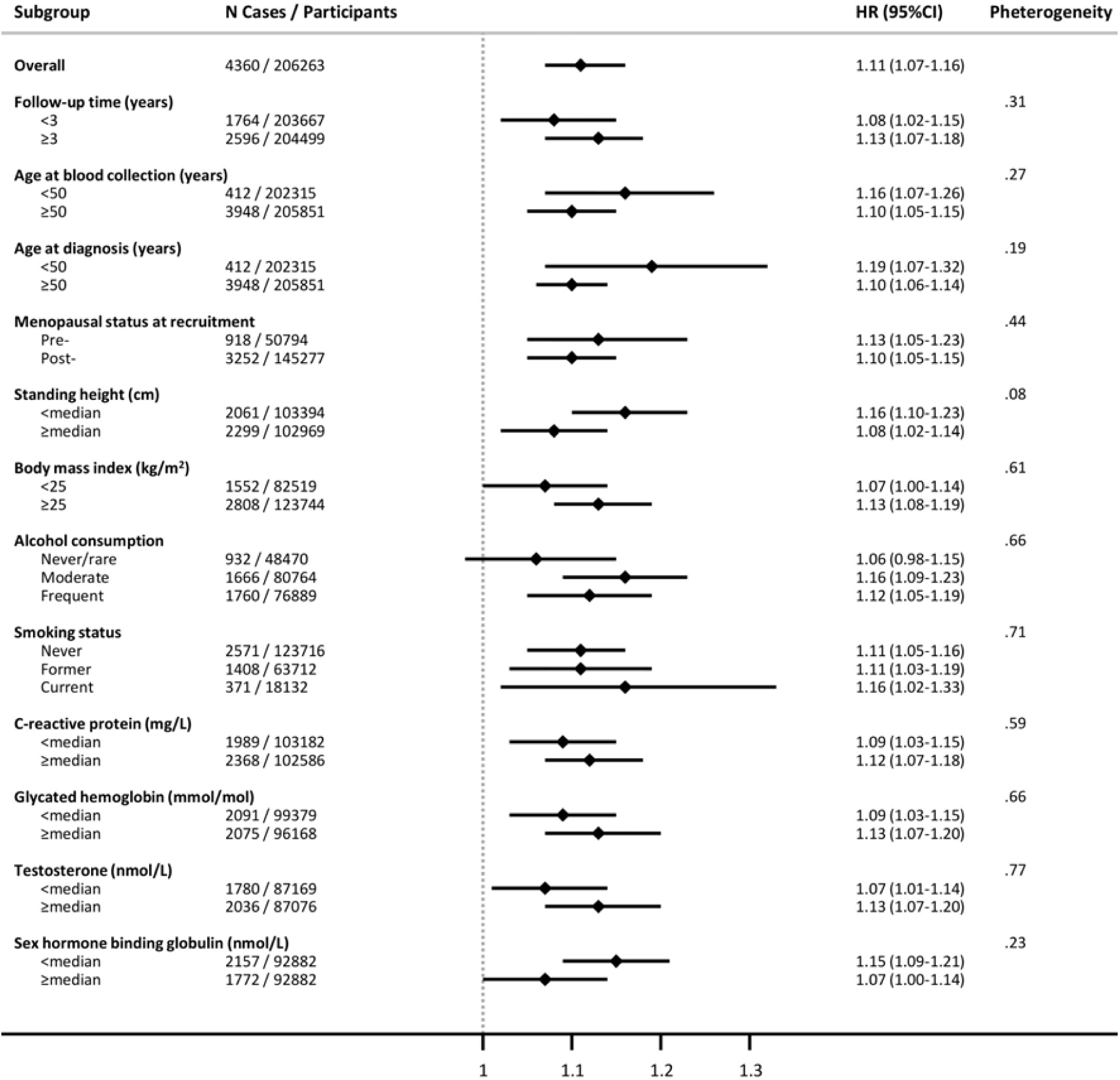
Subgroup analyses of the association between circulating insulin-like growth factor-1 (IGF-1) concentrations and breast cancer risk in the UK Biobank (per 5 nmol/L increment) HR=hazard ratio; CI=confidence interval. Multivariable Cox regression model using age as the underlying time variable and stratified by Townsend deprivation index (fifths), region of the recruitment assessment center, and age at recruitment (5-year categories). Models adjusted for total physical activity (<10, 10-<20, 20-<40, 40-<60, ≥60 MET hours per week, unknown); height (per 10 cm); alcohol consumption frequency (never, special occasions only, 1-3 times per month, 1-2 times per week, 3-4 times per week, daily/almost daily, unknown); smoking status and intensity (never, former, current- <15 per day, current- ≥15 per day, current-intensity unknown, unknown); educational level (CSEs/O-levels/GCSEs or equivalent, NVQ/HND/HNC/A-levels/AS-levels or equivalent, other professional qualifications, college/university degree, none of the above, unknown); ever use of hormone replacement therapy (no, yes, unknown); parity, age at first birth (nulliparous; 1-2, <25 years; 1-2, 25- 30 years; 1-2, ≥30 years; 1-2, unknown; ≥3, <25 years; ≥3, 25-30 years; ≥3, ≥30 years; ≥3, unknown; unknown); the interaction between menopausal status and body mass index (kg/m^2^); and circulating concentrations (fifths, missing/unknown) of C-reactive protein (CRP; mg/L), glycated hemoglobin (HbA1c; mmol/mol), testosterone (nmol/L), and sex hormone blinding globulin (SHBG; nmol/L). Median values: height = 162 cm; C-reactive protein (CRP) = 1.3 mg/L; glycated hemoglobin (HbA1c) = 35.1 mmol/mol; testosterone = 1 nmol/L; sex hormone binding globulin (SHBG) = 56.3 nmol/L. HRs per 5 nmol/L increment were additionally corrected for regression dilution using a regression dilution ratio (0.74) obtained from the subsample of women with repeat IGF-1 measurements.

### Mendelian randomization

#### Association between genetically-predicted circulating insulin-like growth factor 1 (IGF-1) concentrations and breast cancer risk

In the inverse-variance weighted model, a 5 nmol/L increment in genetically-predicted IGF-1 concentrations was associated with greater breast cancer risk (OR=1.05, 95%CI=1.01-1.10; Pvalue=0.02) (Table 3). IGF-1 was positively associated with ER+ (OR=1.06, 95%CI=1.01-1.11; Pvalue=0.03), but not ER^-^ (OR=1.02, 95%CI=0.96-1.08; Pvalue=0.58) tumors (Pheterogeneity=0.32). Evidence of effect heterogeneity (Cochran’s Q Pvalues <0.0001) was found for all models. Little evidence of directional pleiotropy for the breast cancer and ER+ breast cancer models (MR-Egger intercept Pvalues>0.2) was found, with similar magnitude effect estimates observed for the weighted median and lower powered MR-Egger models (Table 3). A similar pattern of results was found when the MR-PRESSO test detected outlier SNPs were excluded from the models (Table S4). The funnel plots for the IGF-1 instrument indicated a symmetric distribution of effect estimates (Figure S1).

**Table 3.** Mendelian randomization estimates between circulating concentrations of insulin-like growth factor-1 (IGF-1) and insulin-like growth factor-binding protein-3 (IGFBP-3) and risk of breast cancer (N=122,977 breast cancer cases and N=105,974 controls)

| Mendelian randomization |  |  |  |
| --- | --- | --- | --- |
|  | Inverse-variance-weighted (IVW) | Weighted median | MR-Egger |
|  | OR (95%CI)<br>per 5 nmol/L<br>increment | OR (95%CI)<br>per 5 nmol/L<br>increment | OR (95%CI)<br>per 5 nmol/L<br>increment |
| **Insulin-like growth factor-1 (IGF-1) (N=265 SNPs)** |  |  |  |
| Breast cancer | 1.05 (1.01-1.10) | 1.08 (1.03-1.13) | 1.12 (1.01-1.25) |
| Breast cancer, ER<sup>+</sup> | 1.06 (1.01-1.11) | 1.10 (1.04-1.18) | 1.12 (1.00-1.26) |
| Breast cancer, ER<sup>-</sup> | 1.02 (0.96-1.08) | 1.03 (0.95-1.11) | 1.17 (1.02-1.34) |
|  | Inverse-variance-weighted (IVW) | Weighted median | MR-Egger |
|  | OR (95%CI)<br>per 1-SD increment | OR (95%CI)<br>per 1-SD increment | OR (95%CI)<br>per 1-SD increment |
| **Insulin-like growth factor-binding protein-3 (IGFBP-3) (N=4 SNPs)** |  |  |  |
| Breast cancer | 1.00 (0.97-1.04) | 1.00 (0.96-1.04) | 0.95 (0.87-1.03) |
| Breast cancer, ER<sup>+</sup> | 1.00 (0.96-1.05) | 1.00 (0.95-1.04) | 0.92 (0.83-1.02) |
| Breast cancer, ER<sup>-</sup> | 0.98 (0.92-1.04) | 0.98 (0.91-1.05) | 1.03 (0.88-1.20) |

Abbreviations: ER=estrogen receptor; OR=odds ratio; CI=confidence interval; SD=standard deviation.

#### Association between genetically-predicted circulating insulin-like growth factor-binding protein-3 (IGFBP-3) concentrations and breast cancer risk

No association was found between genetically-predicted circulating IGFBP-3 concentrations and breast cancer risk (Table 3). A similar null result was found for the weighted median, MR-Egger, and leave–one–out sensitivity analyses (Table 3, Table S5).

## Discussion

In the observational analyses of UK Biobank data, we found that higher circulating concentrations of IGF-1 were associated with greater breast cancer risk. This relationship was consistent for pre- and post-menopausal women and by follow-up time. Consistent with this finding, in our MR analyses, we found a positive association between genetically-predicted IGF-1 concentrations and breast cancer risk, with this effect restricted to ER+ tumors. These results support a probable causal role of the IGF pathway in ER+ breast cancer development.

The positive association found between circulating IGF-1 concentrations and breast cancer in our UK Biobank observational analyses was monotonic and consistent with the result from the Endogenous Hormones and Breast Cancer Collaborative Group, a pooled analysis that included 4,790 breast cancers cases and 9,428 matched controls from 17 prospective studies^8^. Similar to the pooled analysis, this positive association did not differ by menopausal status, follow-up time, and subgroups of other breast cancer risk factors. Our analysis, which included 4,360 incident breast cancer cases, is the largest single study to examine the IGF-1 and breast cancer relationship. Uniquely, measurements of circulating IGF-1 and other biomarkers were available in the full UK Biobank cohort, and we were able to adjust our multivariable models for other serologic factors related to both circulating IGF-1 concentrations and breast cancer risk, namely testosterone, SHBG, CRP, and HbA1c^23-27^. The risk estimates for the IGF-1 and breast cancer relationship were largely unchanged after multivariable statistical adjustment for these biomarkers and other established risk factors. A further unique aspect of our analysis was our correction for regression dilution bias using the repeat IGF-1 measurements available for 6,711 women, thereby mitigating the combined effects of measurement error and within-person variability on our risk estimates^9^. This correction resulted in a strengthening of the positive association, supporting the likelihood that previous epidemiological studies that relied on a single measurement of IGF-1 concentrations underestimated the strength of the positive association with breast cancer risk. For instance, the Endogenous Hormones and Breast Cancer Collaborative Group reported an OR of 1.25 (95%CI=1.13-1.39) for an 80 percentile difference in IGF-1^8^; while the HR at an equivalent scale in the current UK Biobank analysis rose from 1.26 (95%CI=1.15-1.38) to 1.37 (95%CI=1.21-1.55) after correction for regression dilution bias.

Observational analyses may be subject to residual confounding and reverse causality, making causal inference challenging. We conducted MR analyses of the associations between IGF-1, IGFBP-3 and breast cancer risk. MR uses germline genetic variants as proxies (instrumental variables) to allow causal inference between a given exposure and outcome. Unlike traditional observational analyses, MR analyses should be largely free of confounding and reverse causality due to the random assortment of alleles at meiosis and germline genetic variants being fixed at conception, and thus unaffected by the disease process. For IGF-1, the MR analysis yielded a positive effect estimate similar to our observational analysis. This positive effect was only present for ER+ and not ER^-^ breast cancer, a result consistent with earlier observational^8^ and laboratory evidence^28^, which suggests that crosstalk from estrogen signaling pathways may influence the IGF-1 and breast cancer relationship.

The bioavailability of IGF-1 in circulation is partly regulated by IGFBPs, with most bound to IGFBP-3. In addition to enhancing or inhibiting actions of IGF ligands, *in vitro* experimental models suggest that IGFBP-3 can inhibit breast cancer proliferation and induce apoptosis^4,29^. Our MR analyses found no evidence of an association between genetically-predicted IGFBP-3 concentrations and breast cancer risk. This result is consistent with the Endogenous Hormones and Breast Cancer Collaborative Group analysis that reported a null association for IGFBP-3 concentrations and breast cancer risk after the multivariable models were adjusted for circulating IGF-1 concentrations^8^. Taken together, these results provide little evidence of IGFBP-3 having a direct effect, independent of its role in IGF ligand binding, in breast cancer development.

A fundamental assumption of MR analyses is that the genetic instrument should not influence the outcome via a different biological pathway from the exposure of interest (horizontal pleiotropy). We conducted various sensitivity analyses to assess the possible influence of horizontal pleiotropy on our causal estimates, and our results were robust to these various tests. The possibility exists, however, that our results may have been influenced by pleiotropy from other unmeasured IGF axis components^30^. To date, the only GWAS analyses that have been conducted for components of this pathway are for IGF-1 and IGFBP-3, therefore the extent of this possible pleiotropy within the IGF axis is uncertain. Genetic instruments are now required for other IGF system components to disentangle possible biological effects of specific ligands and binding proteins in breast cancer development.

The current study is the largest and most comprehensive investigation of the role of IGF-1 in breast cancer development. A limitation of our observational analysis is that tumor subtype data are currently unavailable in the UK Biobank; however, these data were available for our MR analyses and we found that the positive effect for IGF-1 and breast cancer was only present for ER+ tumors. For our MR analyses, we were unable to stratify the analyses by menopausal status; however, our observational analyses found no difference in the IGF-1 and breast cancer relationship between pre- and post-menopausal women. Our use of summary-level data for our MR analyses meant that we were unable to examine possible non-linear effects or whether the associations between IGF-1 and breast cancer differed according to subgroups of other risk factors (e.g. BMI, alcohol consumption); however, in our observational analyses, which yielded a similar association to our MR result, we found a linear association between IGF-1 and breast cancer and detected no heterogeneity by subgroups of other risk factors.

In conclusion, given that plausible biological mechanisms have been identified^1,2^, our observational and MR results support a probable causal relationship between circulating IGF-1 concentrations and breast cancer. This result suggests that pharmacological or lifestyle interventions targeting the IGF pathway may be beneficial in preventing breast tumorigenesis.

## Acknowledgements

This work has been conducted using the UK Biobank Resource under Application Numbers 3248 and 25897 and we express our gratitude to the participants and those involved in building the resource. UK Biobank is an open access resource. Bona fide researchers can apply to use the UK Biobank dataset by registering and applying at http://ukbiobank.ac.uk/register-apply/.

The breast cancer genome-wide association analyses were supported by the Government of Canada through Genome Canada and the Canadian Institutes of Health Research, the ‘Ministère de l’Économie, de la Science et de l’Innovation du Québec’ through Genome Québec and grant PSR-SIIRI-701, The National Institutes of Health (U19 CA148065, X01HG007492), Cancer Research UK (C1287/A10118, C1287/A16563, C1287/A10710) and The European Union (HEALTH-F2-2009-223175 and H2020 633784 and 634935). All studies and funders are listed in Michailidou et al^10^.

AK is supported by the Wellcome Trust (LEAP 205212/Z/16/Z). GF, APC, RCT and TJK are supported by Cancer Research UK (C8221/A19170).

RMM is supported by a Cancer Research UK program grant (C18281/A19169) and the National Institute for Health Research (NIHR) Bristol Biomedical Research Centre. The views expressed are those of the author(s) and not necessarily those of the NIHR or the Department of Health and Social Care.

