## Supplementary Materials for "Insulin-like growth factor-1 (IGF-1), insulin-like growth factor-binding protein-3 (IGFBP-3) and breast cancer risk: observational and Mendelian randomization analyses"

**Table S1. Summary information on the insulin-like growth factor-1 (IGF-1) and insulin-like growth factor-binding protein-3 (IGFBP-3) genetic instruments used in the Mendelian randomization analyses**

**Table S2. Association parameters of instrumental SNPs used in the insulin-like growth factor-1 (IGF-1) and insulin-like growth factor-binding protein-3 (IGFBP-3) genetic instruments**

**Table S3. Subgroup analyses of the association between circulating insulin-like growth factor-1 (IGF-1) concentrations and breast cancer risk in the UK Biobank**

**Table S4. Mendelian randomization estimates between circulating insulin-like growth factor-1 (IGF-1) and risk of breast cancer, a sensitivity analysis excluding outlying SNPs detected by MR-PRESSO**

**Table S5. Mendelian randomization leave-one-out analysis for insulin-like growth factor-binding protein-3 (IGFBP-3) concentrations and risk of breast cancer**

**Figure S1. Funnel plots of risk estimates of insulin-like growth factor-1 (IGF-1) and (A) breast cancer, (B) breast cancer ER^+^, (C) and breast cancer ER^-^, against instrumental strength**

| **Table S1. Summary information on the insulin-like growth factor-1 (IGF-1) and insulin-like growth factor-binding protein-3 (IGFBP-3) genetic instruments used in the Mendelian randomization analyses** | | | | |
| --- | --- | --- | --- | --- |
| **Exposure** | **Source/Publication** | ***N* SNPs** | **Variance explained (%)** | **Mean (SD)** |
| Insulin-like growth factor-1 (IGF-1) | UK Biobank/Neale Lab 2019^18^  (estimates for women only) | 265 | 5.2 | 21.0 (5.8) nmol/L |
| Insulin-like growth factor-binding protein-3 (IGFBP-3) | Meta-analysis/Teumer et al 2016^30^  (estimates for men and women) | 4 | 6.1 | 3340.4 (863.3) ng/mL |
| *N* SNPs=Number of SNPs; SD=standard deviation. | | | | |

| **Table S2. Association parameters of instrumental SNPs used in the insulin-like growth factor-1 (IGF-1) and insulin-like growth factor-binding protein-3 (IGFBP-3) genetic instruments** | | | | | | | | | | | | | | | |
| --- | --- | --- | --- | --- | --- | --- | --- | --- | --- | --- | --- | --- | --- | --- | --- |
|  |  |  |  |  | | **Association parameters for IGF-1 concentration** | | | **Association parameters for breast cancer** | | | **Association parameters for breast cancer ER^+ve^** | | **Association parameters for breast cancer ER^-ve^** | |
| **SNP** | **Chr** | **Position** | **Effect allele** | **Other allele** | | **effect** | | **se** | **effect** | | **se** | **effect** | **se** | **effect** | **se** |
| **Insulin-like growth factor-1 (IGF-1) (N=265 SNPs)** | | | | | | | | | | | | | | | |
| rs903908 | 1 | 2202967 | C | T | | 0.105 | 0.018 | | | -0.010 | 0.007 | -0.009 | 0.008 | -0.002 | 0.012 |
| rs111656050 | 1 | 26464923 | A | G | | 0.186 | 0.024 | | | 0.000 | 0.008 | 0.006 | 0.010 | -0.007 | 0.015 |
| rs282178 | 1 | 26899687 | A | C | | 0.130 | 0.024 | | | 0.014 | 0.008 | 0.017 | 0.010 | 0.009 | 0.015 |
| rs193084249 | 1 | 26987646 | A | G | | 0.535 | 0.060 | | | -0.029 | 0.021 | -0.044 | 0.025 | -0.048 | 0.038 |
| rs1042114 | 1 | 29138975 | G | T | | 0.145 | 0.026 | | | -0.016 | 0.009 | -0.017 | 0.011 | -0.002 | 0.017 |
| rs605709 | 1 | 44058467 | T | C | | 0.162 | 0.019 | | | -0.006 | 0.006 | 0.003 | 0.008 | -0.022 | 0.012 |
| rs4314918 | 1 | 44377503 | A | G | | 0.148 | 0.024 | | | 0.013 | 0.008 | 0.015 | 0.010 | -0.014 | 0.015 |
| rs1046011 | 1 | 65898996 | T | C | | 0.135 | 0.020 | | | 0.002 | 0.007 | -0.008 | 0.008 | 0.012 | 0.012 |
| rs78511209 | 1 | 91414243 | G | C | | 0.193 | 0.031 | | | 0.018 | 0.011 | 0.024 | 0.013 | 0.003 | 0.020 |
| rs469721 | 1 | 91530001 | T | C | | 0.335 | 0.023 | | | -0.009 | 0.008 | -0.005 | 0.010 | 0.006 | 0.015 |
| rs1977658 | 1 | 107607037 | T | G | | 0.113 | 0.019 | | | 0.001 | 0.007 | -0.001 | 0.008 | 0.019 | 0.012 |
| rs4970836 | 1 | 109821797 | G | A | | 0.155 | 0.022 | | | -0.006 | 0.008 | 0.001 | 0.009 | -0.003 | 0.014 |
| rs1127313 | 1 | 154556425 | G | A | | 0.133 | 0.018 | | | 0.000 | 0.006 | 0.002 | 0.008 | -0.009 | 0.012 |
| rs77369503 | 1 | 163027266 | G | A | | 0.277 | 0.050 | | | -0.004 | 0.020 | 0.004 | 0.024 | -0.057 | 0.036 |
| rs12749024 | 1 | 176522365 | T | C | | 0.433 | 0.026 | | | 0.010 | 0.009 | 0.009 | 0.011 | 0.007 | 0.017 |
| rs2152318 | 1 | 179293511 | T | C | | 0.140 | 0.021 | | | 0.010 | 0.007 | 0.017 | 0.009 | -0.006 | 0.013 |
| rs11118991 | 1 | 208300128 | C | T | | 0.147 | 0.021 | | | -0.001 | 0.007 | -0.001 | 0.009 | -0.005 | 0.014 |
| rs7541039 | 1 | 214176779 | T | C | | 0.118 | 0.021 | | | 0.023 | 0.008 | 0.024 | 0.009 | 0.033 | 0.014 |
| rs17597773 | 1 | 221054761 | C | G | | 0.218 | 0.021 | | | -0.003 | 0.008 | -0.001 | 0.009 | 0.002 | 0.014 |
| rs7517340 | 1 | 243710190 | C | T | | 0.164 | 0.024 | | | 0.004 | 0.008 | 0.008 | 0.010 | 0.019 | 0.015 |
| rs28707636 | 2 | 25935577 | A | G | | 0.294 | 0.021 | | | 0.006 | 0.008 | 0.008 | 0.009 | -0.009 | 0.014 |
| rs4665972 | 2 | 27598097 | C | T | | 0.379 | 0.019 | | | 0.009 | 0.006 | 0.009 | 0.008 | 0.008 | 0.012 |
| rs145957642 | 2 | 27806793 | G | GT | | 0.126 | 0.023 | | | 0.017 | 0.008 | 0.017 | 0.010 | 0.012 | 0.015 |
| rs9309054 | 2 | 40598041 | A | T | | 0.105 | 0.018 | | | -0.004 | 0.006 | -0.001 | 0.007 | -0.006 | 0.011 |
| rs78481334 | 2 | 43796135 | A | T | | 0.150 | 0.027 | | | 0.024 | 0.010 | 0.026 | 0.011 | 0.034 | 0.017 |
| rs2459981 | 2 | 58127925 | A | C | | 0.104 | 0.019 | | | -0.011 | 0.006 | -0.012 | 0.008 | -0.011 | 0.012 |
| rs12471768 | 2 | 64928603 | C | T | | 0.144 | 0.020 | | | 0.019 | 0.007 | 0.023 | 0.009 | 0.016 | 0.013 |
| rs11545482 | 2 | 70315987 | C | T | | 0.582 | 0.064 | | | 0.024 | 0.026 | 0.034 | 0.031 | 0.049 | 0.050 |
| rs11123170 | 2 | 113978940 | G | C | | 0.118 | 0.019 | | | 0.001 | 0.007 | 0.001 | 0.008 | 0.001 | 0.012 |
| rs17400325 | 2 | 178565913 | C | T | | 0.311 | 0.046 | | | 0.011 | 0.016 | 0.014 | 0.019 | 0.006 | 0.029 |
| rs57267144 | 2 | 203392479 | G | C | | 0.140 | 0.021 | | | -0.011 | 0.007 | -0.007 | 0.009 | -0.030 | 0.013 |
| rs646295 | 2 | 219408558 | C | A | | 0.117 | 0.018 | | | -0.003 | 0.006 | -0.007 | 0.008 | -0.009 | 0.012 |
| rs2607748 | 3 | 14158725 | C | T | | 0.104 | 0.018 | | | -0.008 | 0.006 | -0.007 | 0.008 | -0.003 | 0.012 |
| rs11928797 | 3 | 33457493 | A | C | | 0.161 | 0.028 | | | 0.005 | 0.011 | 0.004 | 0.013 | 0.018 | 0.019 |
| rs4683324 | 3 | 47251284 | C | T | | 0.107 | 0.018 | | | 0.006 | 0.006 | 0.019 | 0.008 | -0.017 | 0.011 |
| rs76979176 | 3 | 51264296 | A | G | | 0.331 | 0.060 | | | 0.049 | 0.021 | 0.037 | 0.026 | 0.056 | 0.040 |
| rs7628689 | 3 | 88216647 | G | A | | 0.158 | 0.025 | | | -0.027 | 0.009 | -0.023 | 0.010 | -0.049 | 0.016 |
| rs3772102 | 3 | 98502628 | G | T | | 0.123 | 0.018 | | | 0.007 | 0.006 | 0.006 | 0.007 | -0.006 | 0.011 |
| rs6764700 | 3 | 101486050 | A | G | | 0.200 | 0.035 | | | 0.025 | 0.012 | 0.032 | 0.015 | 0.014 | 0.023 |
| rs7639292 | 3 | 107295665 | C | T | | 0.142 | 0.024 | | | 0.002 | 0.008 | 0.004 | 0.010 | -0.012 | 0.015 |
| rs687339 | 3 | 135932359 | T | C | | 0.237 | 0.022 | | | -0.014 | 0.007 | -0.010 | 0.009 | -0.028 | 0.014 |
| rs68088905 | 3 | 138849533 | G | C | | 0.115 | 0.020 | | | -0.004 | 0.007 | -0.005 | 0.009 | 0.005 | 0.014 |
| rs1582874 | 3 | 141115219 | T | C | | 0.187 | 0.018 | | | -0.047 | 0.006 | -0.055 | 0.007 | -0.016 | 0.011 |
| rs1345410 | 3 | 170291615 | C | T | | 0.138 | 0.025 | | | 0.016 | 0.008 | 0.014 | 0.010 | 0.011 | 0.015 |
| rs5398 | 3 | 170715830 | A | G | | 0.113 | 0.020 | | | 0.011 | 0.007 | 0.000 | 0.009 | 0.044 | 0.013 |
| rs149846151 | 3 | 172123761 | GGA | G | | 0.261 | 0.019 | | | 0.006 | 0.007 | 0.012 | 0.008 | -0.003 | 0.013 |
| rs9879333 | 3 | 186381220 | G | A | | 0.128 | 0.021 | | | 0.013 | 0.007 | 0.013 | 0.009 | 0.013 | 0.013 |
| rs114303452 | 4 | 3449915 | A | G | | 0.552 | 0.088 | | | 0.027 | 0.030 | 0.048 | 0.035 | 0.034 | 0.054 |
| rs4234797 | 4 | 7219907 | A | G | | 0.236 | 0.019 | | | 0.004 | 0.006 | 0.009 | 0.008 | -0.017 | 0.012 |
| rs12499617 | 4 | 7736890 | G | A | | 0.106 | 0.018 | | | 0.002 | 0.007 | 0.007 | 0.008 | -0.001 | 0.013 |
| rs3069978 | 4 | 39696922 | A | G | | 0.133 | 0.020 | | | 0.004 | 0.007 | -0.006 | 0.009 | 0.001 | 0.014 |
| rs7689003 | 4 | 45125825 | A | G | | 0.172 | 0.019 | | | 0.013 | 0.007 | 0.021 | 0.008 | -0.006 | 0.013 |
| rs62302688 | 4 | 46448465 | G | A | | 0.184 | 0.031 | | | 0.003 | 0.012 | -0.001 | 0.015 | 0.027 | 0.022 |
| rs34327193 | 4 | 69324066 | T | TA | | 0.169 | 0.020 | | | 0.008 | 0.008 | 0.004 | 0.009 | -0.004 | 0.015 |
| rs989075 | 4 | 69533221 | A | T | | 0.120 | 0.018 | | | -0.010 | 0.007 | -0.008 | 0.008 | -0.020 | 0.013 |
| rs1379935 | 4 | 89751291 | A | G | | 0.154 | 0.028 | | | -0.005 | 0.010 | -0.002 | 0.012 | -0.004 | 0.018 |
| rs6822348 | 4 | 100053894 | T | A | | 0.181 | 0.020 | | | 0.011 | 0.007 | 0.016 | 0.008 | -0.007 | 0.012 |
| rs1229984 | 4 | 100239319 | T | C | | 0.363 | 0.062 | | | 0.011 | 0.017 | 0.020 | 0.020 | 0.041 | 0.029 |
| rs17429745 | 4 | 106038169 | G | T | | 0.171 | 0.020 | | | -0.020 | 0.007 | -0.023 | 0.008 | -0.017 | 0.012 |
| rs111516782 | 4 | 121723035 | TATATA | T | | 0.126 | 0.020 | | | -0.007 | 0.007 | 0.000 | 0.008 | -0.011 | 0.013 |
| rs2054381 | 4 | 178944245 | A | C | | 0.118 | 0.020 | | | 0.000 | 0.007 | -0.004 | 0.008 | -0.003 | 0.013 |
| rs60701 | 5 | 10733776 | T | C | | 0.114 | 0.021 | | | -0.007 | 0.007 | -0.006 | 0.009 | -0.016 | 0.013 |
| rs4703511 | 5 | 35210629 | G | A | | 0.312 | 0.047 | | | 0.001 | 0.017 | 0.020 | 0.020 | -0.020 | 0.030 |
| rs2303808 | 5 | 39074296 | G | C | | 0.126 | 0.018 | | | 0.001 | 0.006 | -0.003 | 0.007 | 0.009 | 0.011 |
| rs55681913 | 5 | 42687629 | C | T | | 0.346 | 0.030 | | | 0.016 | 0.011 | 0.027 | 0.013 | -0.009 | 0.021 |
| rs34315422 | 5 | 43096271 | T | C | | 0.123 | 0.018 | | | -0.012 | 0.007 | -0.003 | 0.008 | -0.016 | 0.012 |
| rs112838464 | 5 | 53308819 | A | G | | 0.163 | 0.029 | | | -0.003 | 0.010 | 0.003 | 0.012 | -0.017 | 0.018 |
| rs40270 | 5 | 55804552 | C | A | | 0.124 | 0.022 | | | 0.009 | 0.008 | 0.006 | 0.009 | 0.029 | 0.014 |
| rs151196451 | 5 | 59028725 | T | TTAAGTGACTTTCACATGGA | | 0.218 | 0.020 | | | 0.028 | 0.007 | 0.028 | 0.008 | 0.039 | 0.013 |
| rs475058 | 5 | 78341978 | C | T | | 0.106 | 0.018 | | | 0.002 | 0.006 | 0.001 | 0.007 | -0.001 | 0.011 |
| rs6452875 | 5 | 89466182 | A | T | | 0.140 | 0.023 | | | 0.002 | 0.008 | -0.004 | 0.009 | 0.001 | 0.014 |
| rs12520928 | 5 | 132305891 | C | A | | 0.236 | 0.024 | | | 0.025 | 0.009 | 0.034 | 0.010 | 0.018 | 0.016 |
| rs12659034 | 5 | 137773525 | T | C | | 0.214 | 0.023 | | | 0.007 | 0.008 | 0.009 | 0.009 | -0.003 | 0.014 |
| rs2974438 | 5 | 168250903 | G | A | | 0.297 | 0.022 | | | 0.000 | 0.008 | -0.005 | 0.010 | 0.002 | 0.015 |
| rs730551 | 6 | 6971753 | C | T | | 0.100 | 0.018 | | | 0.004 | 0.006 | 0.003 | 0.008 | -0.001 | 0.012 |
| rs111553730 | 6 | 25992022 | G | GTATC | | 0.179 | 0.020 | | | 0.007 | 0.007 | 0.009 | 0.008 | 0.014 | 0.013 |
| rs11445433 | 6 | 26474985 | CT | C | | 0.127 | 0.020 | | | 0.024 | 0.007 | 0.024 | 0.008 | 0.024 | 0.013 |
| rs9265824 | 6 | 31311213 | A | T | | 0.139 | 0.020 | | | -0.003 | 0.007 | 0.002 | 0.008 | -0.003 | 0.013 |
| rs3129763 | 6 | 32590925 | G | A | | 0.187 | 0.022 | | | -0.001 | 0.008 | 0.001 | 0.009 | -0.016 | 0.014 |
| rs28532217 | 6 | 32623887 | T | C | | 0.210 | 0.028 | | | 0.008 | 0.010 | 0.010 | 0.011 | 0.011 | 0.018 |
| rs1187115 | 6 | 34172055 | A | C | | 0.147 | 0.024 | | | 0.017 | 0.008 | 0.009 | 0.010 | 0.015 | 0.015 |
| rs9472040 | 6 | 43348512 | A | G | | 0.109 | 0.019 | | | -0.009 | 0.007 | -0.005 | 0.008 | -0.028 | 0.012 |
| rs11967262 | 6 | 43760327 | G | C | | 0.117 | 0.018 | | | -0.025 | 0.006 | -0.028 | 0.008 | -0.014 | 0.012 |
| rs4147614 | 6 | 52658191 | A | G | | 0.132 | 0.019 | | | -0.002 | 0.007 | 0.001 | 0.008 | 0.020 | 0.012 |
| rs949846 | 6 | 79817596 | C | T | | 0.111 | 0.019 | | | 0.010 | 0.006 | 0.017 | 0.008 | 0.003 | 0.012 |
| rs6940544 | 6 | 88000581 | G | A | | 0.142 | 0.018 | | | 0.001 | 0.006 | 0.003 | 0.007 | -0.012 | 0.011 |
| rs7739044 | 6 | 100239365 | C | T | | 0.130 | 0.018 | | | 0.010 | 0.006 | 0.008 | 0.008 | 0.011 | 0.012 |
| rs2764264 | 6 | 108934461 | T | C | | 0.259 | 0.020 | | | 0.003 | 0.007 | 0.000 | 0.008 | -0.014 | 0.012 |
| rs4897175 | 6 | 126657472 | G | A | | 0.244 | 0.018 | | | 0.010 | 0.006 | 0.009 | 0.008 | 0.009 | 0.012 |
| rs12524625 | 6 | 127418688 | A | G | | 0.126 | 0.019 | | | -0.017 | 0.007 | -0.014 | 0.008 | -0.026 | 0.013 |
| rs556493 | 6 | 147549297 | A | G | | 0.117 | 0.018 | | | -0.009 | 0.007 | -0.018 | 0.008 | -0.006 | 0.012 |
| rs2077647 | 6 | 152129077 | C | T | | 0.122 | 0.018 | | | -0.010 | 0.006 | -0.009 | 0.007 | -0.023 | 0.011 |
| rs655370 | 6 | 153413460 | C | G | | 0.114 | 0.018 | | | -0.002 | 0.006 | 0.003 | 0.007 | -0.016 | 0.011 |
| rs112201728 | 6 | 160551486 | T | C | | 0.296 | 0.036 | | | -0.015 | 0.012 | -0.025 | 0.014 | -0.005 | 0.021 |
| rs662138 | 6 | 160564476 | G | C | | 0.154 | 0.023 | | | 0.000 | 0.008 | 0.002 | 0.010 | -0.010 | 0.015 |
| rs3127580 | 6 | 160710851 | T | C | | 0.194 | 0.025 | | | -0.001 | 0.009 | 0.002 | 0.011 | -0.001 | 0.016 |
| rs520829 | 6 | 160767905 | T | G | | 0.185 | 0.018 | | | -0.006 | 0.006 | -0.011 | 0.007 | -0.001 | 0.011 |
| rs34205994 | 6 | 166316105 | GC | G | | 0.206 | 0.019 | | | 0.020 | 0.007 | 0.010 | 0.008 | 0.039 | 0.013 |
| rs111957895 | 7 | 880983 | G | C | | 0.155 | 0.024 | | | 0.016 | 0.009 | 0.019 | 0.011 | 0.022 | 0.016 |
| rs12699547 | 7 | 2015970 | C | T | | 0.113 | 0.019 | | | 0.018 | 0.006 | 0.022 | 0.008 | 0.001 | 0.012 |
| rs34495733 | 7 | 6763641 | G | A | | 0.230 | 0.026 | | | -0.004 | 0.009 | -0.001 | 0.011 | -0.012 | 0.017 |
| rs4719393 | 7 | 14219213 | T | G | | 0.133 | 0.020 | | | -0.005 | 0.007 | -0.009 | 0.009 | -0.001 | 0.013 |
| rs1079866 | 7 | 41470093 | G | C | | 0.178 | 0.027 | | | -0.011 | 0.009 | -0.011 | 0.011 | 0.005 | 0.016 |
| rs117244063 | 7 | 45758261 | G | A | | 0.449 | 0.069 | | | 0.001 | 0.024 | 0.007 | 0.029 | 0.041 | 0.044 |
| rs2854746 | 7 | 45960645 | G | C | | 0.255 | 0.019 | | | 0.004 | 0.006 | 0.004 | 0.008 | 0.006 | 0.012 |
| rs260359 | 7 | 46467020 | A | C | | 0.245 | 0.039 | | | -0.037 | 0.014 | -0.031 | 0.017 | -0.047 | 0.026 |
| rs10230290 | 7 | 46471388 | G | A | | 0.378 | 0.028 | | | 0.020 | 0.010 | 0.014 | 0.012 | 0.046 | 0.018 |
| rs112993474 | 7 | 46618570 | T | C | | 0.411 | 0.069 | | | -0.017 | 0.032 | 0.011 | 0.037 | -0.093 | 0.057 |
| rs62452750 | 7 | 46702781 | T | C | | 0.268 | 0.019 | | | 0.010 | 0.007 | 0.020 | 0.009 | -0.016 | 0.013 |
| rs2116643 | 7 | 46718310 | C | G | | 0.346 | 0.019 | | | 0.008 | 0.007 | 0.011 | 0.008 | 0.009 | 0.012 |
| rs17172759 | 7 | 46781660 | A | C | | 0.474 | 0.087 | | | 0.007 | 0.033 | 0.007 | 0.038 | -0.039 | 0.058 |
| rs11764457 | 7 | 46951010 | C | T | | 0.312 | 0.040 | | | -0.001 | 0.015 | 0.002 | 0.017 | 0.017 | 0.026 |
| rs13232120 | 7 | 72983310 | T | A | | 0.284 | 0.027 | | | 0.034 | 0.011 | 0.031 | 0.013 | 0.029 | 0.019 |
| rs7792321 | 7 | 108070955 | G | C | | 0.115 | 0.019 | | | 0.002 | 0.007 | -0.002 | 0.008 | -0.012 | 0.012 |
| rs13246909 | 7 | 113753054 | T | C | | 0.102 | 0.019 | | | 0.005 | 0.007 | 0.006 | 0.008 | -0.008 | 0.012 |
| rs157928 | 7 | 130581358 | C | T | | 0.255 | 0.019 | | | 0.001 | 0.007 | -0.004 | 0.008 | -0.021 | 0.013 |
| rs273956 | 7 | 137603188 | A | G | | 0.111 | 0.018 | | | -0.005 | 0.007 | -0.008 | 0.008 | -0.008 | 0.012 |
| rs7789908 | 7 | 150530196 | G | T | | 0.125 | 0.022 | | | -0.002 | 0.008 | -0.006 | 0.010 | 0.012 | 0.015 |
| rs41341748 | 8 | 16012594 | G | A | | 0.526 | 0.086 | | | 0.010 | 0.033 | 0.015 | 0.038 | 0.023 | 0.061 |
| rs7842117 | 8 | 59313897 | C | G | | 0.114 | 0.018 | | | 0.016 | 0.006 | 0.017 | 0.008 | 0.008 | 0.012 |
| rs3214527 | 8 | 81400281 | C | CAG | | 0.112 | 0.020 | | | -0.012 | 0.007 | -0.011 | 0.008 | -0.055 | 0.013 |
| rs2737205 | 8 | 116610180 | T | C | | 0.112 | 0.018 | | | 0.008 | 0.006 | 0.006 | 0.008 | -0.002 | 0.011 |
| rs7830798 | 8 | 134609410 | G | A | | 0.153 | 0.023 | | | 0.006 | 0.008 | 0.016 | 0.009 | -0.001 | 0.014 |
| rs7012213 | 8 | 135660469 | T | A | | 0.107 | 0.020 | | | -0.020 | 0.007 | -0.022 | 0.008 | -0.011 | 0.012 |
| rs456179 | 9 | 4851966 | T | C | | 0.262 | 0.028 | | | 0.007 | 0.010 | 0.013 | 0.012 | -0.036 | 0.018 |
| rs12552790 | 9 | 4864237 | C | T | | 0.289 | 0.043 | | | -0.022 | 0.018 | -0.038 | 0.022 | 0.027 | 0.033 |
| rs72701648 | 9 | 5149890 | C | A | | 0.292 | 0.042 | | | 0.039 | 0.017 | 0.026 | 0.020 | 0.083 | 0.031 |
| rs10757291 | 9 | 22161884 | G | A | | 0.123 | 0.018 | | | 0.003 | 0.006 | 0.013 | 0.007 | -0.017 | 0.011 |
| rs10992863 | 9 | 96445803 | A | G | | 0.131 | 0.020 | | | 0.001 | 0.007 | -0.003 | 0.009 | -0.007 | 0.013 |
| rs28393820 | 9 | 98281825 | T | C | | 0.110 | 0.019 | | | -0.014 | 0.007 | -0.018 | 0.008 | 0.000 | 0.012 |
| rs78509281 | 9 | 109566543 | T | C | | 0.285 | 0.042 | | | 0.017 | 0.015 | 0.012 | 0.018 | 0.016 | 0.028 |
| rs5900219 | 9 | 119312168 | TATTAAAAGTA | T | | 0.170 | 0.026 | | | 0.040 | 0.009 | 0.032 | 0.010 | 0.048 | 0.016 |
| rs2416964 | 9 | 128360413 | C | T | | 0.118 | 0.019 | | | 0.001 | 0.007 | 0.004 | 0.008 | -0.010 | 0.012 |
| rs61854729 | 10 | 5232942 | C | T | | 0.326 | 0.024 | | | 0.023 | 0.008 | 0.035 | 0.010 | 0.008 | 0.015 |
| rs2050905 | 10 | 22027644 | A | G | | 0.127 | 0.019 | | | -0.034 | 0.006 | -0.049 | 0.008 | 0.023 | 0.012 |
| rs2068888 | 10 | 94839642 | G | A | | 0.121 | 0.018 | | | 0.012 | 0.006 | 0.012 | 0.007 | -0.003 | 0.011 |
| rs12219199 | 10 | 95332517 | T | C | | 0.246 | 0.043 | | | 0.025 | 0.015 | 0.032 | 0.017 | 0.053 | 0.026 |
| rs116454156 | 10 | 95347041 | A | G | | 0.545 | 0.074 | | | 0.046 | 0.025 | 0.046 | 0.029 | 0.082 | 0.045 |
| rs2274224 | 10 | 96039597 | G | C | | 0.106 | 0.018 | | | 0.006 | 0.006 | 0.011 | 0.008 | -0.006 | 0.012 |
| rs11190741 | 10 | 102655313 | C | T | | 0.160 | 0.018 | | | -0.019 | 0.006 | -0.014 | 0.008 | -0.017 | 0.012 |
| rs4917985 | 10 | 104624072 | G | A | | 0.104 | 0.019 | | | -0.012 | 0.006 | -0.013 | 0.008 | -0.006 | 0.012 |
| rs3858325 | 10 | 117988795 | T | C | | 0.145 | 0.018 | | | -0.001 | 0.006 | -0.001 | 0.008 | -0.007 | 0.012 |
| rs112945586 | 11 | 1608445 | C | T | | 0.199 | 0.034 | | | 0.000 | 0.011 | -0.001 | 0.014 | 0.000 | 0.021 |
| rs3858525 | 11 | 2058421 | G | C | | 0.128 | 0.018 | | | -0.005 | 0.006 | -0.010 | 0.008 | 0.017 | 0.011 |
| rs10840340 | 11 | 2116492 | G | C | | 0.360 | 0.019 | | | 0.002 | 0.007 | -0.005 | 0.008 | 0.015 | 0.012 |
| rs116862756 | 11 | 2123710 | G | A | | 0.448 | 0.063 | | | -0.029 | 0.026 | -0.023 | 0.030 | -0.010 | 0.049 |
| rs17885785 | 11 | 2167850 | T | C | | 0.421 | 0.023 | | | -0.006 | 0.008 | -0.005 | 0.009 | -0.008 | 0.014 |
| rs78396389 | 11 | 2209620 | C | T | | 0.482 | 0.048 | | | -0.023 | 0.016 | -0.014 | 0.019 | 0.014 | 0.029 |
| rs11024614 | 11 | 18326758 | C | T | | 0.112 | 0.019 | | | 0.014 | 0.007 | 0.013 | 0.008 | -0.008 | 0.012 |
| rs7127069 | 11 | 28325412 | G | A | | 0.138 | 0.020 | | | 0.001 | 0.007 | 0.004 | 0.009 | 0.011 | 0.013 |
| rs10767832 | 11 | 30199020 | T | C | | 0.119 | 0.019 | | | 0.012 | 0.007 | 0.018 | 0.008 | -0.006 | 0.012 |
| rs3136447 | 11 | 46744368 | C | T | | 0.137 | 0.022 | | | 0.003 | 0.007 | 0.000 | 0.009 | 0.010 | 0.014 |
| rs7479361 | 11 | 48109793 | G | A | | 0.217 | 0.019 | | | 0.007 | 0.007 | 0.000 | 0.008 | 0.012 | 0.012 |
| rs10657263 | 11 | 49690460 | C | G | | 0.129 | 0.018 | | | 0.010 | 0.006 | 0.004 | 0.008 | 0.013 | 0.012 |
| rs12790261 | 11 | 66988048 | A | C | | 0.190 | 0.033 | | | -0.012 | 0.014 | -0.006 | 0.017 | 0.005 | 0.027 |
| rs4980662 | 11 | 69307011 | C | G | | 0.101 | 0.018 | | | -0.028 | 0.006 | -0.041 | 0.007 | 0.005 | 0.011 |
| rs643090 | 11 | 118567115 | C | T | | 0.105 | 0.019 | | | 0.011 | 0.007 | 0.012 | 0.008 | 0.021 | 0.013 |
| rs11218882 | 11 | 122771664 | C | T | | 0.109 | 0.019 | | | -0.004 | 0.007 | 0.006 | 0.008 | -0.010 | 0.013 |
| rs11054429 | 12 | 11868537 | G | C | | 0.158 | 0.019 | | | 0.016 | 0.007 | 0.019 | 0.008 | 0.014 | 0.012 |
| rs10841648 | 12 | 20954557 | A | C | | 0.121 | 0.019 | | | -0.008 | 0.007 | -0.007 | 0.008 | -0.003 | 0.012 |
| rs9738365 | 12 | 31997635 | A | C | | 0.267 | 0.021 | | | 0.012 | 0.007 | 0.011 | 0.009 | 0.002 | 0.013 |
| rs1259768 | 12 | 32074187 | C | G | | 0.229 | 0.029 | | | -0.005 | 0.011 | -0.004 | 0.013 | -0.012 | 0.020 |
| rs11425044 | 12 | 56488735 | C | CA | | 0.109 | 0.019 | | | 0.000 | 0.007 | 0.002 | 0.008 | 0.008 | 0.013 |
| rs324019 | 12 | 57486647 | A | G | | 0.102 | 0.019 | | | 0.005 | 0.007 | 0.009 | 0.008 | 0.008 | 0.012 |
| rs7484541 | 12 | 57714803 | A | T | | 0.124 | 0.022 | | | -0.015 | 0.008 | -0.013 | 0.010 | -0.007 | 0.014 |
| rs1351394 | 12 | 66351826 | C | T | | 0.130 | 0.018 | | | 0.005 | 0.006 | 0.001 | 0.007 | 0.011 | 0.011 |
| rs3741675 | 12 | 94108529 | A | G | | 0.106 | 0.018 | | | 0.009 | 0.006 | 0.007 | 0.007 | 0.007 | 0.011 |
| rs10860237 | 12 | 98157010 | A | G | | 0.173 | 0.020 | | | 0.008 | 0.007 | 0.010 | 0.008 | -0.004 | 0.012 |
| rs79685578 | 12 | 102231032 | C | T | | 0.190 | 0.034 | | | -0.007 | 0.015 | -0.010 | 0.018 | -0.028 | 0.028 |
| rs6539022 | 12 | 102353197 | G | T | | 0.181 | 0.029 | | | -0.020 | 0.011 | -0.018 | 0.013 | -0.017 | 0.020 |
| rs4567531 | 12 | 102399379 | C | A | | 0.317 | 0.024 | | | -0.004 | 0.008 | -0.010 | 0.010 | 0.000 | 0.015 |
| rs117688359 | 12 | 102403033 | G | A | | 0.444 | 0.058 | | | 0.033 | 0.021 | 0.038 | 0.025 | 0.031 | 0.039 |
| rs117403498 | 12 | 102567644 | A | G | | 0.511 | 0.065 | | | 0.029 | 0.024 | 0.017 | 0.029 | 0.039 | 0.044 |
| rs186537413 | 12 | 102828079 | C | T | | 0.369 | 0.047 | | | -0.037 | 0.019 | -0.028 | 0.023 | -0.066 | 0.035 |
| rs10860867 | 12 | 102841826 | T | C | | 0.879 | 0.074 | | | -0.046 | 0.025 | -0.057 | 0.029 | -0.033 | 0.047 |
| rs10860878 | 12 | 102963550 | C | T | | 0.130 | 0.018 | | | -0.011 | 0.007 | -0.014 | 0.008 | -0.002 | 0.013 |
| rs4764939 | 12 | 103522952 | T | C | | 0.149 | 0.018 | | | 0.003 | 0.006 | 0.007 | 0.007 | -0.001 | 0.011 |
| rs11830764 | 12 | 111515020 | C | G | | 0.384 | 0.036 | | | 0.013 | 0.014 | 0.011 | 0.016 | 0.021 | 0.024 |
| rs3741698 | 12 | 115109223 | G | C | | 0.132 | 0.020 | | | 0.041 | 0.007 | 0.053 | 0.008 | 0.015 | 0.013 |
| rs12810788 | 12 | 116196322 | A | G | | 0.151 | 0.023 | | | 0.010 | 0.008 | 0.002 | 0.010 | 0.018 | 0.015 |
| rs3742039 | 12 | 120649970 | C | T | | 0.102 | 0.018 | | | -0.007 | 0.006 | -0.007 | 0.008 | -0.007 | 0.012 |
| rs679833 | 12 | 121220820 | G | A | | 0.160 | 0.020 | | | -0.004 | 0.007 | -0.003 | 0.008 | -0.020 | 0.013 |
| rs1800574 | 12 | 121416864 | T | C | | 0.695 | 0.054 | | | 0.001 | 0.020 | -0.019 | 0.023 | 0.048 | 0.035 |
| rs3213547 | 12 | 121435181 | A | G | | 0.277 | 0.039 | | | -0.017 | 0.014 | -0.015 | 0.016 | -0.022 | 0.025 |
| rs76750172 | 13 | 28395297 | T | C | | 0.329 | 0.059 | | | -0.020 | 0.025 | -0.022 | 0.029 | -0.031 | 0.045 |
| rs79212157 | 13 | 40766836 | T | C | | 0.186 | 0.023 | | | 0.032 | 0.008 | 0.037 | 0.010 | 0.029 | 0.014 |
| rs1170155 | 13 | 42702711 | T | C | | 0.117 | 0.019 | | | -0.001 | 0.007 | -0.004 | 0.008 | 0.005 | 0.012 |
| rs4942553 | 13 | 47155975 | G | A | | 0.113 | 0.020 | | | -0.006 | 0.008 | -0.003 | 0.009 | -0.021 | 0.014 |
| rs9583151 | 13 | 107666257 | C | T | | 0.101 | 0.018 | | | 0.012 | 0.007 | 0.014 | 0.008 | 0.011 | 0.012 |
| rs1755770 | 14 | 36666760 | T | C | | 0.141 | 0.019 | | | 0.008 | 0.007 | 0.010 | 0.008 | -0.010 | 0.012 |
| rs2093210 | 14 | 60957279 | C | T | | 0.134 | 0.019 | | | 0.002 | 0.006 | 0.001 | 0.008 | 0.013 | 0.012 |
| rs168961 | 14 | 69282930 | G | A | | 0.101 | 0.018 | | | 0.006 | 0.006 | 0.004 | 0.008 | 0.008 | 0.012 |
| rs8020365 | 14 | 79937216 | A | T | | 0.121 | 0.022 | | | 0.013 | 0.008 | 0.015 | 0.009 | 0.010 | 0.014 |
| rs10142298 | 14 | 93909792 | T | C | | 0.135 | 0.020 | | | 0.001 | 0.007 | -0.004 | 0.008 | 0.001 | 0.012 |
| rs10136874 | 14 | 101202022 | G | T | | 0.110 | 0.018 | | | 0.012 | 0.006 | 0.015 | 0.007 | 0.008 | 0.011 |
| rs11070390 | 15 | 43428110 | A | T | | 0.103 | 0.019 | | | -0.017 | 0.007 | -0.014 | 0.008 | -0.018 | 0.012 |
| rs201393666 | 15 | 43677979 | C | A | | 0.805 | 0.056 | | | -0.007 | 0.019 | 0.002 | 0.023 | 0.022 | 0.035 |
| rs2414098 | 15 | 51537806 | T | C | | 0.130 | 0.019 | | | -0.018 | 0.006 | -0.028 | 0.008 | -0.003 | 0.012 |
| rs79391862 | 15 | 53739426 | C | A | | 0.425 | 0.078 | | | 0.019 | 0.032 | 0.041 | 0.037 | 0.031 | 0.060 |
| rs17205463 | 15 | 62381413 | C | T | | 0.221 | 0.018 | | | -0.011 | 0.007 | -0.005 | 0.008 | -0.025 | 0.012 |
| rs28434383 | 15 | 67448181 | C | T | | 0.113 | 0.020 | | | -0.004 | 0.007 | 0.003 | 0.008 | -0.031 | 0.012 |
| rs143717852 | 15 | 68382383 | A | G | | 0.224 | 0.038 | | | 0.013 | 0.015 | 0.014 | 0.017 | -0.008 | 0.027 |
| rs5742915 | 15 | 74336633 | C | T | | 0.106 | 0.018 | | | -0.013 | 0.006 | -0.012 | 0.007 | -0.017 | 0.011 |
| rs1994147 | 15 | 95829524 | A | T | | 0.138 | 0.020 | | | -0.011 | 0.008 | -0.009 | 0.010 | -0.021 | 0.015 |
| rs8034564 | 15 | 99190601 | A | G | | 0.149 | 0.018 | | | -0.002 | 0.007 | -0.009 | 0.008 | -0.005 | 0.012 |
| rs2715439 | 15 | 99492313 | C | T | | 0.117 | 0.018 | | | -0.014 | 0.006 | -0.013 | 0.008 | -0.026 | 0.012 |
| rs12924916 | 16 | 950224 | C | T | | 0.124 | 0.020 | | | -0.004 | 0.007 | -0.009 | 0.008 | 0.000 | 0.013 |
| rs76819935 | 16 | 1110581 | T | C | | 0.526 | 0.040 | | | -0.007 | 0.020 | -0.011 | 0.024 | 0.028 | 0.037 |
| rs7187586 | 16 | 1146109 | G | C | | 0.106 | 0.018 | | | -0.007 | 0.008 | -0.013 | 0.009 | 0.006 | 0.014 |
| rs8048693 | 16 | 1811565 | A | G | | 0.286 | 0.019 | | | 0.001 | 0.007 | 0.006 | 0.008 | -0.008 | 0.012 |
| rs7197536 | 16 | 1846241 | T | C | | 0.223 | 0.026 | | | 0.007 | 0.010 | 0.004 | 0.012 | 0.017 | 0.018 |
| rs11076826 | 16 | 4427010 | G | C | | 0.132 | 0.021 | | | -0.014 | 0.008 | -0.019 | 0.010 | 0.003 | 0.015 |
| rs12924995 | 16 | 9028645 | A | C | | 0.107 | 0.018 | | | -0.010 | 0.006 | -0.013 | 0.008 | -0.013 | 0.012 |
| rs35083214 | 16 | 12567217 | C | G | | 0.121 | 0.021 | | | -0.006 | 0.007 | -0.005 | 0.008 | -0.010 | 0.013 |
| rs80275162 | 16 | 28863517 | A | T | | 0.107 | 0.019 | | | 0.005 | 0.006 | 0.009 | 0.008 | -0.005 | 0.012 |
| rs140820592 | 16 | 31021880 | G | T | | 0.180 | 0.019 | | | 0.016 | 0.007 | 0.008 | 0.008 | 0.022 | 0.012 |
| rs72787359 | 16 | 51431230 | A | C | | 0.197 | 0.035 | | | 0.003 | 0.018 | 0.001 | 0.021 | -0.013 | 0.032 |
| rs112941090 | 16 | 69130106 | C | G | | 0.137 | 0.024 | | | -0.005 | 0.008 | -0.009 | 0.010 | -0.023 | 0.015 |
| rs11396409 | 16 | 69560330 | AC | A | | 0.159 | 0.022 | | | -0.009 | 0.008 | -0.009 | 0.009 | -0.024 | 0.015 |
| rs6564889 | 16 | 81591496 | C | T | | 0.130 | 0.018 | | | -0.003 | 0.007 | 0.000 | 0.008 | -0.016 | 0.012 |
| rs67044703 | 16 | 81738525 | C | T | | 0.127 | 0.018 | | | 0.012 | 0.007 | 0.008 | 0.008 | 0.021 | 0.012 |
| rs11149612 | 16 | 83980965 | C | T | | 0.120 | 0.018 | | | 0.002 | 0.007 | 0.006 | 0.009 | -0.009 | 0.013 |
| rs9896759 | 17 | 5465268 | T | C | | 0.106 | 0.018 | | | -0.005 | 0.007 | -0.002 | 0.008 | -0.018 | 0.012 |
| rs12952818 | 17 | 17988591 | G | A | | 0.134 | 0.019 | | | 0.019 | 0.006 | 0.025 | 0.008 | 0.019 | 0.011 |
| rs1045929 | 17 | 38175426 | C | T | | 0.146 | 0.019 | | | 0.008 | 0.006 | 0.002 | 0.008 | 0.009 | 0.012 |
| rs117292219 | 17 | 40451137 | G | T | | 0.196 | 0.032 | | | -0.011 | 0.011 | -0.005 | 0.013 | -0.016 | 0.021 |
| rs142377191 | 17 | 61649170 | A | G | | 0.837 | 0.063 | | | 0.064 | 0.028 | 0.030 | 0.033 | 0.048 | 0.050 |
| rs2854152 | 17 | 61986027 | G | A | | 0.208 | 0.019 | | | 0.007 | 0.007 | 0.006 | 0.008 | -0.004 | 0.012 |
| rs3760237 | 17 | 62051110 | C | T | | 0.141 | 0.018 | | | 0.002 | 0.007 | -0.008 | 0.008 | 0.007 | 0.012 |
| rs76708468 | 17 | 62206299 | C | T | | 0.519 | 0.047 | | | 0.040 | 0.023 | 0.063 | 0.027 | -0.021 | 0.043 |
| rs144830625 | 17 | 62601428 | T | C | | 0.401 | 0.063 | | | -0.007 | 0.032 | -0.014 | 0.037 | 0.019 | 0.058 |
| rs8178824 | 17 | 64224775 | T | C | | 0.427 | 0.054 | | | 0.013 | 0.019 | 0.012 | 0.023 | 0.015 | 0.035 |
| rs77542162 | 17 | 67081278 | G | A | | 0.380 | 0.060 | | | 0.003 | 0.024 | 0.010 | 0.029 | -0.003 | 0.046 |
| rs4567812 | 18 | 1646432 | A | G | | 0.185 | 0.020 | | | 0.023 | 0.007 | 0.018 | 0.008 | 0.023 | 0.012 |
| rs12462529 | 19 | 4359525 | G | A | | 0.126 | 0.021 | | | 0.020 | 0.007 | 0.020 | 0.009 | 0.016 | 0.014 |
| rs12462204 | 19 | 4772763 | G | C | | 0.163 | 0.027 | | | 0.006 | 0.010 | 0.006 | 0.012 | 0.019 | 0.018 |
| rs9676789 | 19 | 4973852 | A | C | | 0.201 | 0.019 | | | -0.017 | 0.007 | -0.019 | 0.008 | -0.003 | 0.012 |
| rs11668724 | 19 | 7208526 | G | A | | 0.133 | 0.022 | | | 0.004 | 0.008 | 0.006 | 0.010 | 0.015 | 0.014 |
| rs8105174 | 19 | 10347032 | C | T | | 0.245 | 0.023 | | | -0.016 | 0.009 | -0.017 | 0.010 | -0.012 | 0.016 |
| rs280498 | 19 | 10482699 | A | C | | 0.126 | 0.020 | | | -0.020 | 0.007 | -0.026 | 0.009 | -0.010 | 0.014 |
| rs11085744 | 19 | 10819967 | T | C | | 0.170 | 0.018 | | | -0.014 | 0.007 | -0.022 | 0.008 | -0.007 | 0.012 |
| rs72477287 | 19 | 36735275 | AC | A | | 0.111 | 0.019 | | | 0.002 | 0.007 | 0.002 | 0.008 | -0.021 | 0.013 |
| rs2722722 | 19 | 44832875 | A | G | | 0.103 | 0.019 | | | -0.003 | 0.007 | 0.001 | 0.008 | -0.010 | 0.012 |
| rs142998071 | 19 | 47744170 | T | G | | 0.243 | 0.044 | | | 0.021 | 0.018 | 0.001 | 0.021 | 0.083 | 0.032 |
| rs10425975 | 19 | 48384635 | G | A | | 0.200 | 0.024 | | | 0.017 | 0.008 | 0.019 | 0.010 | 0.009 | 0.015 |
| rs2386994 | 19 | 48463194 | C | G | | 0.158 | 0.024 | | | 0.016 | 0.009 | 0.016 | 0.010 | 0.018 | 0.016 |
| rs739320 | 19 | 49261368 | C | T | | 0.151 | 0.019 | | | -0.001 | 0.007 | 0.004 | 0.009 | -0.004 | 0.013 |
| rs10405357 | 19 | 54759666 | T | C | | 0.102 | 0.018 | | | 0.005 | 0.007 | 0.012 | 0.008 | 0.001 | 0.012 |
| rs2208030 | 20 | 3355567 | T | C | | 0.123 | 0.018 | | | 0.004 | 0.006 | 0.005 | 0.008 | 0.012 | 0.012 |
| rs73125628 | 20 | 20066701 | C | T | | 0.293 | 0.020 | | | 0.005 | 0.007 | 0.009 | 0.009 | -0.007 | 0.013 |
| rs35179160 | 20 | 20369563 | G | GT | | 0.154 | 0.021 | | | 0.010 | 0.008 | 0.007 | 0.010 | 0.017 | 0.015 |
| rs6082358 | 20 | 21230455 | C | T | | 0.145 | 0.019 | | | -0.001 | 0.007 | -0.002 | 0.008 | 0.004 | 0.013 |
| rs6058104 | 20 | 33271052 | A | G | | 0.162 | 0.024 | | | -0.027 | 0.009 | -0.023 | 0.010 | -0.032 | 0.016 |
| rs76126732 | 20 | 42087632 | T | A | | 0.203 | 0.037 | | | 0.000 | 0.013 | -0.001 | 0.016 | 0.041 | 0.024 |
| rs11698822 | 20 | 49066506 | C | G | | 0.216 | 0.036 | | | -0.010 | 0.013 | -0.014 | 0.016 | -0.011 | 0.024 |
| rs185799410 | 20 | 57466093 | G | T | | 0.333 | 0.058 | | | 0.086 | 0.024 | 0.084 | 0.028 | 0.066 | 0.044 |
| rs76484331 | 20 | 62422504 | A | C | | 0.245 | 0.039 | | | -0.012 | 0.015 | -0.001 | 0.017 | -0.016 | 0.027 |
| rs8126001 | 20 | 62711459 | T | C | | 0.144 | 0.018 | | | 0.028 | 0.007 | 0.029 | 0.008 | 0.020 | 0.012 |
| rs113455659 | 21 | 37448312 | AAAC | A | | 0.184 | 0.019 | | | -0.008 | 0.007 | -0.010 | 0.008 | 0.003 | 0.012 |
| rs2000467 | 22 | 24241533 | A | G | | 0.115 | 0.019 | | | -0.016 | 0.007 | -0.009 | 0.008 | -0.025 | 0.012 |
| rs8138950 | 22 | 29448643 | C | T | | 0.107 | 0.018 | | | -0.005 | 0.006 | -0.007 | 0.008 | -0.007 | 0.011 |
| rs5755950 | 22 | 36183988 | T | C | | 0.186 | 0.031 | | | 0.001 | 0.012 | 0.005 | 0.014 | -0.001 | 0.022 |
| rs6519133 | 22 | 39096602 | T | C | | 0.174 | 0.019 | | | 0.015 | 0.007 | 0.016 | 0.008 | 0.014 | 0.012 |
| rs200579451 | 22 | 41429704 | TA | T | | 0.101 | 0.019 | | | 0.014 | 0.007 | 0.008 | 0.008 | 0.020 | 0.012 |
| rs9611567 | 22 | 41769754 | A | G | | 0.192 | 0.021 | | | 0.003 | 0.007 | -0.008 | 0.009 | 0.032 | 0.014 |
| **Insulin-like growth factor-binding protein-3 (IGFBP-3) (N=4 SNPs)** | | | | | | | | | | | | | | | |
| rs4234798 | 4 | 7219933 | G | T | 0.095 | | | 0.011 | 0.005 | | 0.006 | 0.009 | 0.008 | -0.017 | 0.012 |
| rs11977526 | 7 | 46008110 | A | G | 0.287 | | | 0.011 | -0.006 | | 0.006 | -0.007 | 0.008 | -0.006 | 0.012 |
| rs700753 | 7 | 46753684 | G | C | 0.158 | | | 0.011 | 0.004 | | 0.007 | 0.004 | 0.008 | 0.005 | 0.012 |
| rs1065656 | 16 | 1838836 | G | C | 0.111 | | | 0.011 | 0.005 | | 0.007 | 0.007 | 0.008 | -0.004 | 0.012 |
| SNP=single nucleotide polymorphism; Chr=chromosome; se=standard error. | | | | | | | | | | | | | | | |

| **Table S3. Subgroup analyses of the association between circulating insulin-like growth factor-1 (IGF-1) concentrations and breast cancer risk in the UK Biobank** | | | | |
| --- | --- | --- | --- | --- |
|  |  |  | **Per 5 nmol/L increment** | **Per 5 nmol/L increment (adjusted) ^a^** |
|  | **N Cases / Participants** | **Pheterogeneity** | **HR (95% CI)** | **HR (95% CI)** |
| **Follow-up time** |  |  |  |  |
| <3 years | 1764 / 203667 | 0.31 | 1.06 (1.02-1.11) | 1.08 (1.02-1.15) |
| ≥3 years | 2596 / 204499 |  | 1.09 (1.05-1.13) | 1.13 (1.07-1.18) |
| **Age at blood collection (years)** |  |  |  |  |
| <50 | 876 / 50270 | 0.27 | 1.11 (1.05-1.18) | 1.16 (1.07-1.26) |
| ≥50 | 3484 / 155993 |  | 1.07 (1.04-1.11) | 1.10 (1.05-1.15) |
| **Age at diagnosis (years)** |  |  |  |  |
| <50 | 412 / 202315 | 0.19 | 1.14 (1.05-1.23) | 1.19 (1.07-1.32) |
| ≥50 | 3948 / 205851 |  | 1.07 (1.04-1.10) | 1.10 (1.06-1.14) |
| **Menopausal status at recruitment** |  |  |  |  |
| Pre- | 918 / 50794 | 0.44 | 1.10 (1.03-1.17) | 1.13 (1.05-1.23) |
| Post- | 3252 / 145277 |  | 1.07 (1.04-1.11) | 1.10 (1.05-1.15) |
| **Standing height** |  |  |  |  |
| <median (<162 cm) | 2061 / 103394 | 0.08 | 1.12 (1.07-1.17) | 1.16 (1.10-1.23) |
| ≥median | 2299 / 102869 |  | 1.06 (1.01-1.10) | 1.08 (1.02-1.14) |
| **Body mass index (kg/m^2^)** |  |  |  |  |
| <25 | 1552 / 82519 | 0.61 | 1.05 (1.00-1.10) | 1.07 (1.00-1.14) |
| ≥25 | 2808 / 123744 |  | 1.10 (1.06-1.14) | 1.13 (1.08-1.19) |
| **Alcohol consumption** |  |  |  |  |
| Never/rare | 932 / 48470 | 0.66 | 1.04 (0.98-1.11) | 1.06 (0.98-1.15) |
| Moderate | 1666 / 80764 |  | 1.11 (1.07-1.17) | 1.16 (1.09-1.23) |
| Frequent | 1760 / 76889 |  | 1.09 (1.04-1.14) | 1.12 (1.05-1.19) |
| **Smoking status** |  |  |  |  |
| Never | 2571 / 123716 | 0.71 | 1.08 (1.04-1.12) | 1.11 (1.05-1.16) |
| Former | 1408 / 63712 |  | 1.08 (1.02-1.13) | 1.11 (1.03-1.19) |
| Current | 371 / 18132 |  | 1.12 (1.01-1.23) | 1.16 (1.02-1.33) |
| **C-reactive protein** |  |  |  |  |
| <median (<1.3 mg/L) | 1989 / 103182 | 0.59 | 1.06 (1.02-1.11) | 1.09 (1.03-1.15) |
| ≥median | 2368 / 102586 |  | 1.09 (1.05-1.13) | 1.12 (1.07-1.18) |
| **Glycated hemoglobin** |  |  |  |  |
| <median (<35.1 mmol/mol) | 2091 / 99379 | 0.66 | 1.06 (1.02-1.11) | 1.09 (1.03-1.15) |
| ≥median | 2075 / 96168 |  | 1.10 (1.05-1.14) | 1.13 (1.07-1.20) |
| **Testosterone** |  |  |  |  |
| <median (<1.0 nmol/L) | 1780 / 87169 | 0.77 | 1.06 (1.01-1.10) | 1.07 (1.01-1.14) |
| ≥median | 2036 / 87076 |  | 1.10 (1.05-1.14) | 1.13 (1.07-1.20) |
| **Sex hormone binding globulin** |  |  |  |  |
| <median (<56.3 nmol/L) | 2157 / 92882 | 0.23 | 1.11 (1.07-1.15) | 1.15 (1.09-1.21) |
| ≥median | 1772 / 92882 |  | 1.05 (1.00-1.10) | 1.07 (1.00-1.14) |
| HR=hazard ratio; CI=confidence interval; SD=standard deviation. | | | | |
| Multivariable Cox regression model using age as the underlying time variable and stratified by Townsend deprivation index (fifths), region of the recruitment assessment center, and age at recruitment (5-year categories). Models adjusted for total physical activity (<10, 10-<20, 20-<40, 40-<60, ≥60 MET hours per week, unknown); height (per 10 cm); alcohol consumption frequency (never, special occasions only, 1-3 times per month, 1-2 times per week, 3-4 times per week, daily/almost daily, unknown); smoking status and intensity (never, former, current- <15 per day, current- ≥15 per day, current- intensity unknown, unknown); educational level (CSEs/O-levels/GCSEs or equivalent, NVQ/HND/HNC/A-levels/AS-levels or equivalent, other professional qualifications, college/university degree, none of the above, unknown); ever use of hormone replacement therapy (no, yes, unknown); parity, age at first birth (nulliparous; 1-2, <25 years; 1-2, 25-30 years; 1-2, ≥30 years; 1-2, unknown; ≥3, <25 years; ≥3, 25-30 years; ≥3, ≥30 years; ≥3, unknown; unknown); the interaction between menopausal status and body mass index (kg/m^2^); and circulating concentrations (fifths, missing/unknown) of C-reactive protein (CRP; mg/L), glycated hemoglobin (HbA1c; mmol/mol), testosterone (nmol/L), and sex hormone blinding globulin (SHBG; nmol/L) | | | | |
| ^a^: HRs per SD increment were additionally corrected for regression dilution using a regression dilution ratio (0.74) obtained from the subsample of women with repeat IGF-1 measurements. | | | | |

| **Table S4. Mendelian randomization estimates between circulating insulin-like growth factor-1 (IGF-1) and risk of breast cancer, a sensitivity analysis excluding outlying SNPs detected by MR-PRESSO** | | | | | | | | | |
| --- | --- | --- | --- | --- | --- | --- | --- | --- | --- |
|  | **Breast cancer ^a^** | | | **Breast cancer, ER^+ b^** | | | **Breast cancer, ER^- c^** | | |
|  | **OR (95% CI)**  **per 5 nmol/L increment** | **Pvalue** | **Pvalue for intercept** | **OR (95% CI)**  **per 5 nmol/L increment** | **Pvalue** | **Pvalue for intercept** | **OR (95% CI)**  **per 5 nmol/L increment** | **Pvalue** | **Pvalue for intercept** |
| Inverse-variance weighted | 1.06 (1.02-1.10) | 0.005 |  | 1.08 (1.04-1.13) | 0.0003 |  | 1.02 (0.97-1.08) | 0.47 |  |
| MR-Egger | 1.11 (1.01-1.22) | 0.03 | 0.31 | 1.11 (1.00-1.22) | 0.06 | 0.67 | 1.16 (1.01-1.32) | 0.03 | 0.04 |
| Weighted median | 1.08 (1.04-1.13) | 0.001 |  | 1.11 (1.05-1.17) | 0.0003 |  | 1.03 (0.95-1.11) | 0.48 |  |
| Abbreviations: ER=estrogen receptor; OR=odds ratio; CI=confidence interval; SD=standard deviation. | | | | | | | | | |
| ^a^: Outlying SNPs excluded (N=8): rs8126001, rs3741698, rs4980662, rs2050905, rs5900219, rs11967262, rs151196451, and rs1582874. | | | | | | | | | |
| ^b^: Outlying SNPs excluded (N=6): rs1582874, rs11967262, rs2050905, rs4980662, rs3741698, and rs2414098. | | | | | | | | | |
| ^c^: Outlying SNP excluded (N=1): rs3214527. | | | | | | | | | |

| **Table S5. Mendelian randomization leave-one-out analysis for insulin-like growth factor-binding protein-3 (IGFBP-3) concentrations and risk of breast cancer** | | |
| --- | --- | --- |
| **SNP excluded** | **Inverse-variance-weighted (IVW)** | **Pvalue** |
|  | **OR (95% CI) per 1-SD increment** |  |
| rs4234798 | 1.00 (0.96-1.03) | 0.81 |
| rs11977526 | 1.04 (0.98-1.10) | 0.25 |
| rs700753 | 0.99 (0.96-1.03) | 0.75 |
| rs1065656 | 1.00 (0.96-1.03) | 0.79 |
| SNP=single nucleotide polymorphism; OR=Odds ratio; CI=confidence interval; SD=standard deviation. | | |

**Figure S1. Funnel plots of risk estimates of insulin-like growth factor-1 (IGF-1) and (A) breast cancer, (B) breast cancer ER^+^, (C) and breast cancer ER^-^, against instrumental strength**

**(A)**

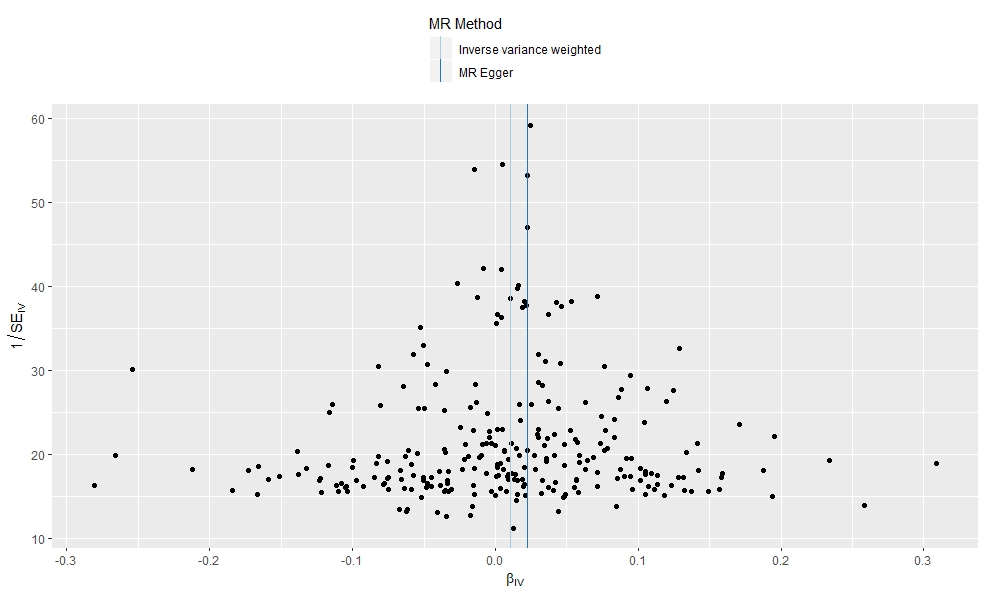

**(B)**

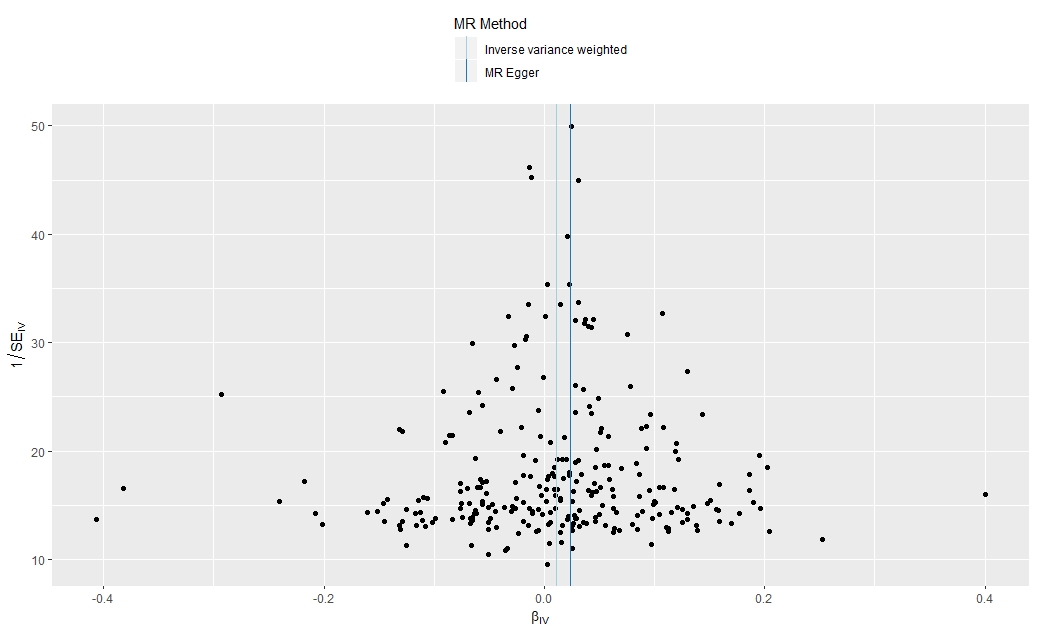

**(C)**

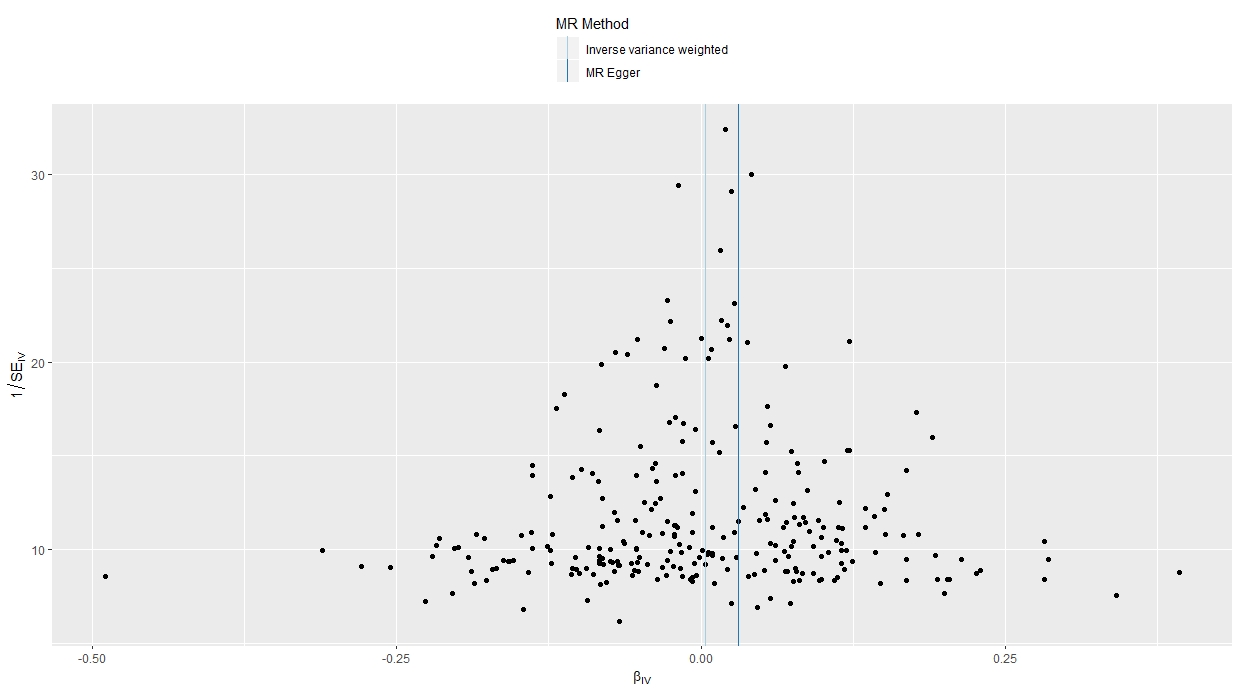

β_IV_ = causal estimate; SE_IV_ = standard error of the instrumental variable
